# Systematic assessment of microenvironment-dependent transcriptional patterns and intercellular communication

**DOI:** 10.1101/2025.01.15.633112

**Authors:** Elena Pareja-Lorente, Patrick Aloy

## Abstract

Understanding cell-cell communication and its dependence on spatial organization is critical for unraveling tissue complexity and organ function. This study integrates single-cell RNA sequencing (scRNA-seq) with spatial transcriptomics (ST) to systematically assess how spatial niches influence gene expression and intercellular communication. Using breast cancer, brain cortex, and heart datasets, our analyses reveal limited global transcriptional changes in cells depending on their spatial microenvironment, with differential gene expression observed in less than half the samples explored. Moreover, cell-cell communication predictions, derived from ligand-receptor pairs, exhibit minimal correlation with spatial colocalization of cell types. Overall, our study underscores the limitations of using scRNA-seq data to capture niche-specific molecular interactions, even when spatial information is leveraged, and it highlights the need for novel strategies to refine our understanding of intercellular communication dynamics at molecular level.

## Introduction

The spatial localization of cells within tissues plays an essential role in determining their biological function^1^, and a complete understanding of the cellular microenvironment and intracellular interactions is fundamental to unravel the complexity of organ function^2^. Indeed, it has been demonstrated that cells are influenced by their microenvironment and neighboring cells, affecting their gene expression and intercellular interactions^3,4^. While single-cell RNA sequencing (scRNA-seq) has enhanced our understanding of gene expression profiles across different cell types, it does not directly capture the spatial arrangement of these cells. On the other hand, spatial transcriptomics (ST) techniques have facilitated the description of gene expression across tissue sections yet lack resolution at the single-cell level. This limitation hampers our ability to detect variations in cellular states based on spatial location and to fully comprehend intercellular communication dynamics^5^, considering the spatial constraints on molecule diffusion from senders to receiver cells^6^.

In the last years, numerous computational methods to infer cell-cell communication have emerged^7^. In the absence of detailed spatial information, most of these approaches rely on the identification of potential receptor-ligand interactions using single-cell sequencing data as a proxy for protein detection (e.g.^8,9^). Indeed, these strategies have identified, for instance, regulatory interactions that prevent harmful immune responses^10^ or unveiled mechanistic insights associated with tissue remodeling in pancreatic tumorigenesis^11^. More recent approaches incorporate spatial transcriptomics to provide additional context to cell-cell communication^6,12^. However, spatial data is scarce and most studies still rely on scRNA-seq data to infer intercellular communication, which has become a standard component of the analytical pipeline^13^. And yet, it is unclear whether the detection of coordinated variations in the expression of receptors and ligands in different cell types is enough to suggest a functional cell-cell communication^14^. Perhaps more importantly, it is key to assess whether a higher number of potential receptor-ligand pairs or a specific transcriptional response translates into a stronger functional interaction between cells^9,15^ or, on the contrary, the identification of a single *bona fide* pair would suffice to ensure an effective communication between cells^16^. Indeed, studies integrating single-cell sequencing with spatial information suggest that widely used methodologies to predict cell-cell interactions are often affected by high rates of false positives^17,18,7^.

In this context, we have developed a systemic approach to investigate single cells in their spatial context. Our methodology leverages existing techniques for assigning spatial coordinates to individual cells using shared embedding spaces or shorter path optimization algorithms. By integrating spatial organization with single-cell gene expression profiles, we defined distinct cellular niches with different cellular compositions, facilitating a deeper examination of cell states influenced by their location and surrounding microenvironment.

## Results and Discussion

### Overview of the methodological strategy

To assess how different cell types might be affected by their surrounding microenvironment (i.e. tissue niche), we have developed a systematic approach to investigate single cells in their spatial context (Figure 1A). By integrating spatial organization with single-cell gene expression profiles, we defined distinct cellular niches with different cellular compositions, facilitating a deeper examination of cell states influenced by their location and tissue niche.

**Figure 1.**
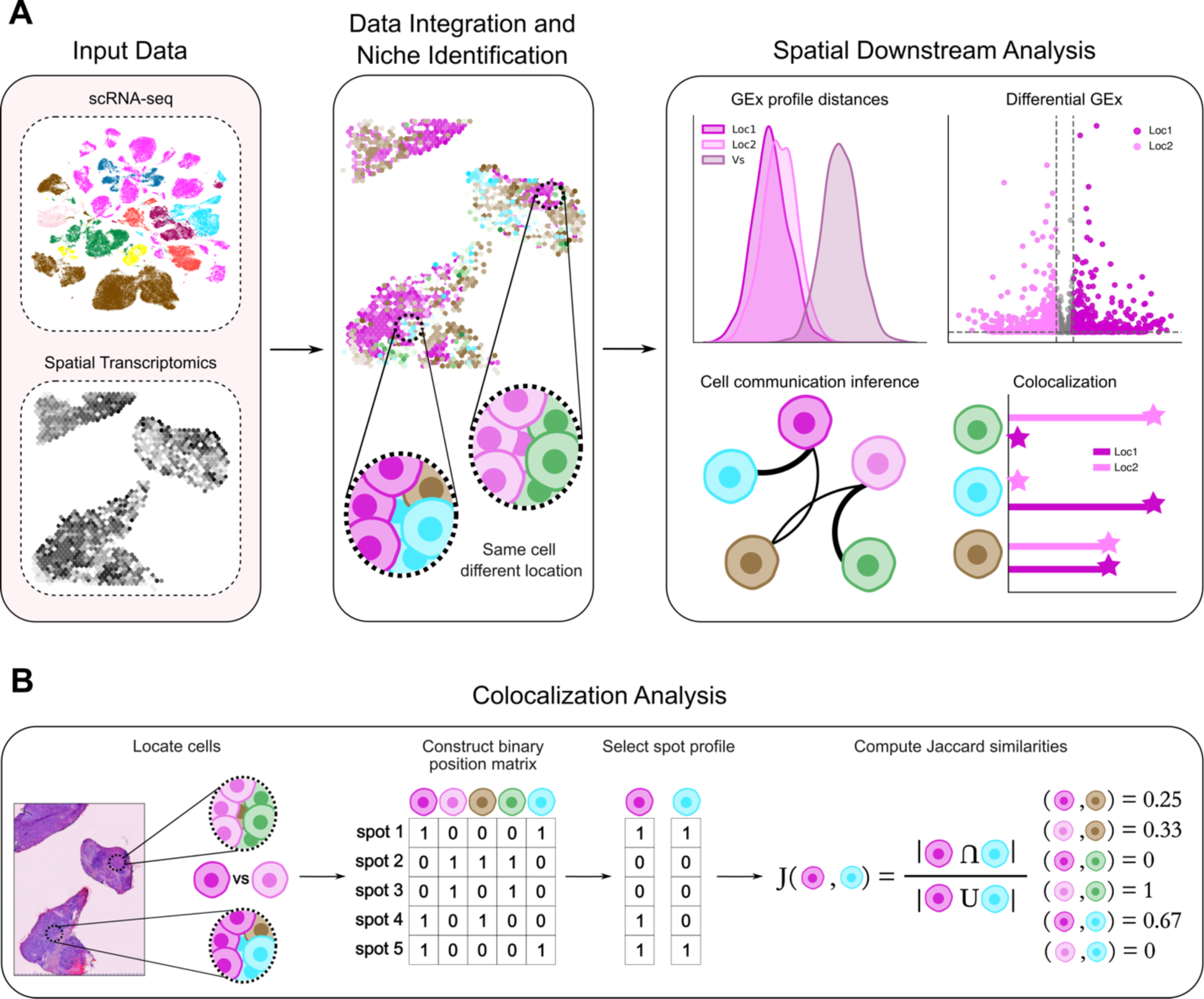
Schematic overview of the pipeline used in the study. **A**. The process begins with the integration of expression matrices and annotations from scRNA-seq and spatial transcriptomics to identify the cellular coalition through different strategies. Following the spatial assignment of single cells, we conduct several downstream analyses to assess the impact of the spatial distribution on cell type dynamics. These downstream analyses include the exploration of differences in Gene Expression profiles (GEx) across different locations, the identification of differentially expressed genes, the investigation of differences in communication dynamics and, finally, the exploration of differences in cellular niches through colocalization analysis. **B**. To quantify the spatial colocalization of different cell types we adapted a previously established strategy^19^. We first constructed a binary position matrix, where a value of 1 indicates the presence of a particular cell type in a spot, and 0 indicates its absence. The columns of this matrix represent the spot profile of the different cell types. To assess the overlap between these position profiles, Jaccard similarities were calculated. This approach allows us to explore differences in cellular colocalization across defined spatial groups.

As first step, we integrate expression matrices and annotations from scRNA-seq and spatial transcriptomics to identify the cellular coalition in each sample slide, and using different strategies. After the assignment of single cells to spatial spots, we conduct several downstream analyses to assess the impact of the spatial distribution on cell type dynamics. In particular, we first investigate the overall differences between gene expression profiles across locations, comparing the distributions of pairwise cosine distances between groups in different niches. We then perform a differential gene expression analysis between groups to identify genes whose expression change depending on their niche. Finally, we examine variations in cell-cell communication dynamics, correlating these with differences in cell-type colocalization analyses (Figure 1B). More specifically, we adapted a recently established strategy to quantify the spatial colocalization of different cell types^19^. We first construct a binary position matrix indicating the presence or absence of a particular cell type in a spot, with the columns of this matrix representing the spot location profile of the different cell types. Then, we calculate Jaccard similarities to assess the overlap between cell-type position profiles, which allows us to explore differences in cellular colocalization across defined spatial groups. We apply our analytical pipeline to three different datasets representing distinct tissues: breast cancer, mouse brain cortex, and heart. A detailed explanation of each step of the analytical pipeline is provided in the *Datasets and Methods* section.

### Case study 1. Exploration of cancer cells in different cellular contexts

Our initial case study comprises four spatial slides from breast cancer patients representing different molecular and anatomical subclasses: two ER^+^ tumors (lobular and ductal, identified as CID4535 and CID4290, respectively) and two TNBC ductal tumors (CID44971 and CID4465)^20^. To integrate patient-specific scRNA-seq measurements within their corresponding ST slide, we annotate the ST spots spatially contextualizing the identified single cells using four different strategies: one based on deconvolution and three on cell mapping techniques (Datasets and Methods). Generally, all four methodologies agree on the cell type distribution across the slides, aligning with pathological annotations of the H&E images provided by the dataset authors (Figure S1). Notably, for the slide from patient CID4535, CytoSPACE^19^ successfully maps T cells to spots in the upper tissue region, designated as “Invasive cancer + lymphocytes” by the authors (Figure 2A), whereas the other three methodologies identify only cancer cells in this area. Additionally, Tangram^21^ assigned very low probabilities to cells in most spots, introducing a level of uncertainty to the mapping results given by this methodology (Figure S1).

**Figure 2.**
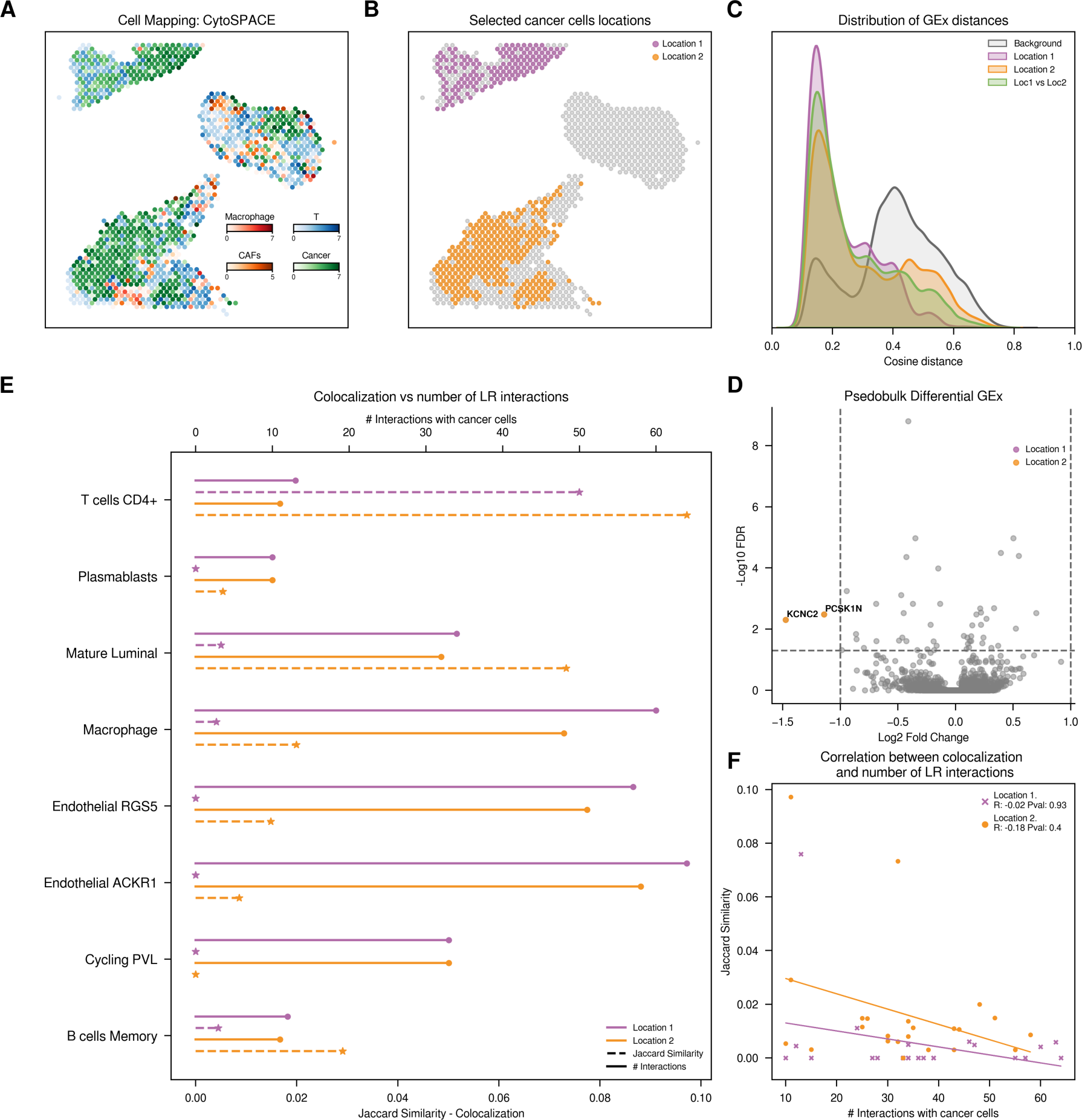
Exploration of cancer cells from CID4535 patient. **A.** Distribution of main cell types across the tissue slide. For visualization purposes, we normalized the CytoSPACE inferred abundances of predominant cell types, with each spatial spot representing the cell type of highest abundance. **B.** Spatial representation of cancer cells according to their assigned location **C.** Distribution of cosine distances between gene expression (GEx) profiles. We analyze the cosine distances between gene expression patterns of cancer cells located within the same or different tissue regions. **D.** Volcano plot with Differential Gene Expression results, highlighting significant upregulated genes in cancer cells of each region. **E.** Comparison of colocalization and cell-cell communication results. Plain line (top axis) represents the count of significant ligand-receptor interactions between each of the defined cancer cells and the y-axis cells. Dotted line (bottom axis) indicates colocalization, measured by the Jaccard similarity index based on the presence or absence of cells within each spot. **F**. Spearman correlation between colocalization and ligand-receptor counts. Color code: Violet cancer cells in location 1, orange cancer cells in location 2.

Overall, we annotated nine cell types on the spatial slides across all patients, including B cells, T cells, cancer epithelial cells, cancer-associated fibroblasts, endothelial cells, myeloid cells, epithelial cells, plasmablast, and perivascular cells (Figures 2 and S2-4). The abundance of these cell types varies by slide, with a particularly low frequency of cancer cells and T cells observed in patient CID4465 (Figure S5). We easily identified distinct spatial regions within the slides, each characterized by different tissue sections on the same spatial slide. In this first case study, we focused on cancer cells to assess how variations in the cellular composition of their spatial niches can affect their gene expression and communication dynamics. Indeed, we could identify significant differences in the cellular environment surrounding cancer cells and define different groups according to their location and different niches (Figures 2B and S2-4B).

To investigate how different cellular environments might globally affect gene expression profiles of specific cells, for each patient, we calculated pairwise cosine distances among the gene expression profiles of all cancer cells within and between spatial groups, and we also compared them to the background of all cell types in the slides (Figure 2C and S2-4C). As expected, we found that the expression profiles of cancer cells are more similar among themselves than when compared to other cell types. However, surprisingly, we also found no differences between cells located in the same niche and those in distinct locations across all ST slides, suggesting that the cellular environment does not significantly impact the overall gene expression of these cells. This result is also supported by UMAP projections of scRNA-seq data, where cancer cells failed to cluster based on their different localizations (Figure S6), and it is in line with previous findings reporting a lack of clustering by tumor cell proximity^19^. The similarity between cancer cells varies by patient, but this heterogeneity is not related to the spatial locations and cellular neighborhood of the cancer cells. To assess whether our global analysis could be masking more subtle but significant gene expression changes, we compared the expression of each gene between the different cancer cell locations (Table S1). As expected, the results of the differential gene expression analyses are quite heterogeneous. For instance, patient CID44971 showed 181 significantly niche-dependent differentially expressed genes (Figure S2D), matching the two different histopathological annotations defined as ductal carcinoma *in situ* (DCIS), where cancer cells are isolated, and invasive cancer with lymphocytes, with cancer cells surrounded by macrophages, T cells and NK cells. Indeed, enrichment analyses on the differentially expressed genes highlighted functional terms associated to immune system regulation (Table S1), suggesting that cancer cells adjacent to lymphocytes trigger defense responses against the immune system ^22^. However, in the other three patients analyzed we found virtually no difference in the expression patterns of cancer cells in different locations, with 2, 0 and 2 differentially expressed genes in CID4535, CID4465 and CID4290 patients, respectively (Figures 2D and S3-4D; Table S1). Note that the number of differentially expressed genes is not related to the expression variability within each location, with average cosine distances ranging from 0.23 for patient CID4535 to 0.62 for patient CID4465 (Figures 2C and S3C).

To investigate the interaction dynamics between spatially defined cancer cells and their cellular niche, we mostly relied on CellPhoneDB^8^, which looks for the coordinate expression of ligands and receptors in paired cell types. In this first case study, CellPhoneDB inferred numerous interactions between cancer cells and other cell types across almost all samples. Specifically, we predicted 1,812 significant interactions in CID4535 patient, 2,648 in CID44971, 280 in CID4465 and 1,387 in CID4290, with interactions between cancer and endothelial cells being the most frequent across all patient samples but CID44971, where CAFs are identified as predominant interactors (Figure S2E). Remarkably, we found a strong correlation in predicted interaction frequency across both groups of spatially defined cancer cells for all samples, with Spearman correlation coefficients of 0.97, 1.0, 0.91 and 0.99, respectively (Figure S7). Such compelling correlations also indicate the absence of detectable differences in the communication patterns between the cancer cell niches, further supporting the observed lack of differences in gene expression patterns between cancer cells in different locations.

Since most ligand-receptor pairs facilitating cell-cell communication act locally, either through physical contacts or limited cytokine diffusion, we next assessed whether the cell types predicted to interact are found in the proximal regions of the tissue slices. To quantify the variance in cellular composition across tissue regions, we labeled spatial spots as 1 if at least one cell of a specific cell type is mapped into that spot, and 0 otherwise. This resulted in a binary matrix where rows represent the spatial spots and the columns the different cell types, describing the spatial distribution of cells in each slice. Using these profiles, we next calculated pairwise Jaccard similarities between cell types to measure the overlap between cancer cells in different locations and the other cell types mapped in the same ST slide (Figure 1B). The low Jaccard similarity scores indicate a tendency for cancer cells to be spatially isolated (i.e. not sharing spots with non-cancer cells), with only 23% of them sharing spatial spots with other cell types. The colocalization analyses revealed differences in the cell composition of the distinct cancer cell locations across all examined ST slides, confirming the presence of different cellular niches across the tissue (Figures 2E and S2-4E). Next, we assessed a potential correlation between colocalization of cancer and other cell types with the number of predicted ligand-receptor interactions, as colocalizing cells should be more likely to interact^23–25^. However, unexpectedly, we observed a total lack of correlation between these two metrics, which, again, was not related to the observed differences in gene expression (Figures 2F and S2-4F). Indeed, we found many instances where cells did not share spatial location but still exhibited elevated counts of predicted ligand-receptor interactions (Figures 2E and S2-4E). For instance, in patient CID4535, cancer cells located in location 1 only colocalize with T cells, whereas cancer cells in location 2 colocalize with various cell types. However, cancer cells in location 1 exhibit high counts of significant predicted LR interactions (940 significant interactions) with other cell types (Figure 2A), questioning the validity of inferred interactions with, for instance, Endothelial ACKR1 cells. Despite the complete lack of spatial co-occurrence with cancer cells in location 1, these cells show the highest count of significant predicted LR interactions with cancer cells in the location (Figure 2E). To evaluate the dependence of our results on the CellPhoneDB strategy and list of ligand-receptor pairs compiled, we repeated our analyses using LIANA^26^, which integrates the ligand-receptor pairs predicted by seven different methods and provides a consensus prediction more accurate than any of the individual methods alone. Again, we observe large discrepancies between the number of predicted interactions between cell types and their spatial colocalization (Figure S8). Indeed, in all the analyzed samples, we find instances of supposedly interacting cell types that are never found in close vicinity. Moreover, we confirmed the reliability of the cell-cell communication results in front of different cell counts by performing subsampling analyses to ensure the same cell numbers across groups before calculating cell-cell communication (Figure S9). Overall, our results show that, despite some variability in the exact number of predicted interactions, the conclusions are robust in different patients and cell-cell communication prediction strategies.

### Case study 2. Exploration of inner and outer layer neurons

To validate the generalization of our findings beyond breast cancer, we extended our analysis to a substantially different tissue: the anterior section of the sagittal mouse brain^27^. Again, as starting point, we integrated scRNA-seq measurements with their corresponding ST slide to spatially contextualize the identified single cells. The mapping results revealed the expected layered structure of the brain, with neurons accurately mapped to their corresponding layers (Figure 3A). Overall, we annotated fifteen different cell types, including neurons from different layers (L2/3, L4, L5 and L6) and major non-neuronal classes (i.e. astrocytes and endothelial cells).

**Figure 3.**
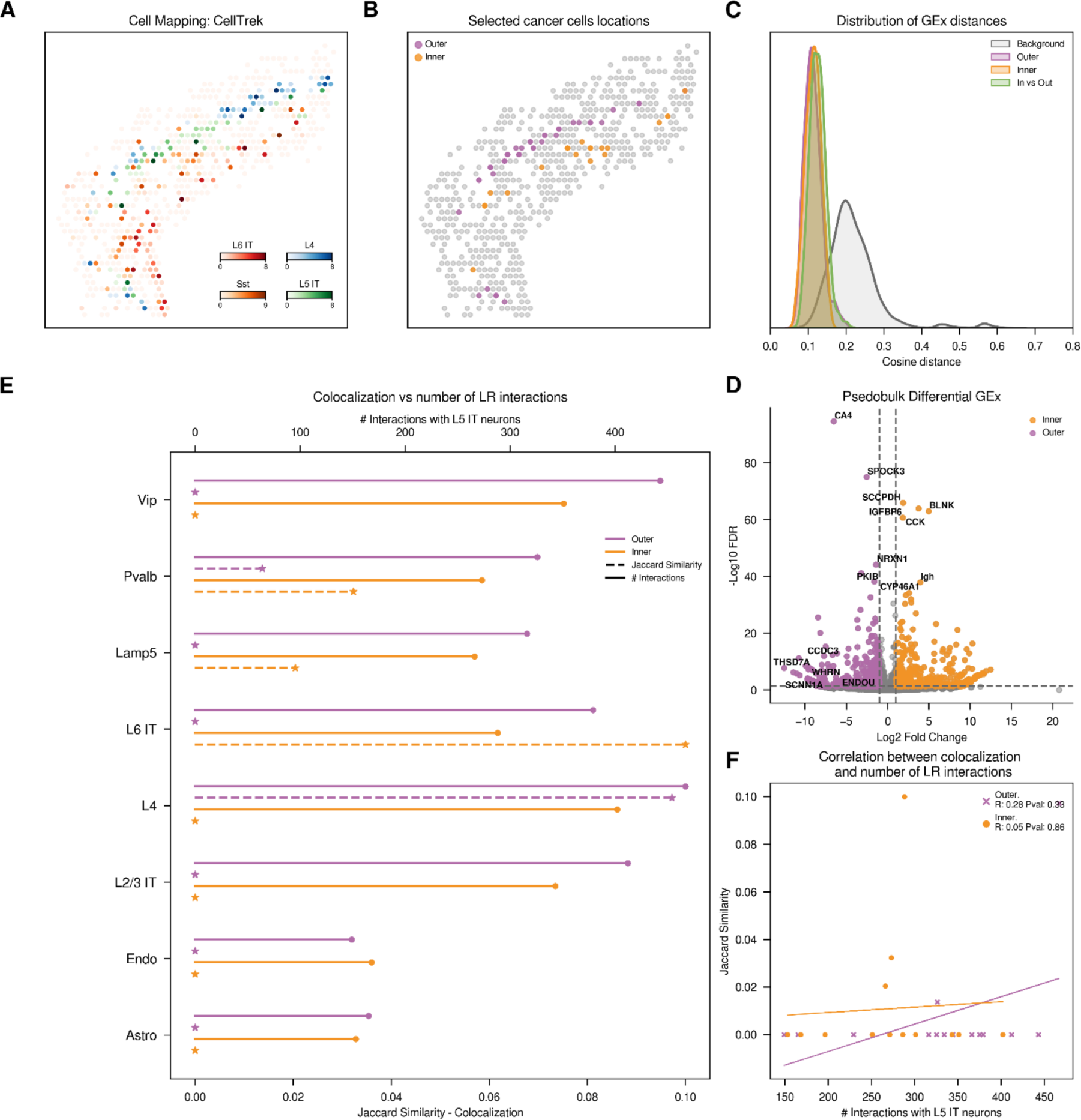
Exploration of L5 IT neurons from mouse brain dataset. **A.** Distribution of adjacent cell types and different neuron types across the tissue slide. For visualization purposes, we normalized the CellTrek inferred abundances of selected cell types, with each spatial spot representing the cell type of highest abundance. **B.** Spatial representation of L5 IT neurons according to their assigned location. **C.** Distribution of cosine distances between gene expression (GEx) profiles. We analyze the cosine distances between gene expression patterns of L5 IT neurons located within the same or different tissue regions (outer or inner part of the L5 layer). **D.** Volcano plot with Differential Gene Expression results, highlighting significant upregulated genes in L5 IT neurons of each region. **E.** Comparison of colocalization and CellPhoneDB cell-cell communication results. Plain line (top axis) represents the count of significant ligand-receptor interactions between each of the defined L5 IT neurons and the y-axis cells. Dotted line (bottom axis) indicates colocalization, measured by the Jaccard similarity index based on the presence or absence of cells within each spot. **F**. Spearman correlation between colocalization and ligand-receptor counts. Color code: Violet neurons in the outer part of the layer, orange neurons in the inner part of the layer.

In this case, to distinguish spatially differentiated groups of cells for subsequent analysis of their expression and communication dynamics, we implemented the Coexp methodology defined in the original CellTrek publication^5^. In brief, the Coexp module identifies spatial gene co-expression modules within a specific cell type. It calculates a spatial distance matrix between each cell of interest and uses it to weight the gene cross-correlation matrix. It then uses the WGCNA approach to identify co-expression modules^28^ and the Seurat AddModuleScore function to assign a cell-level module activity score^29^. In this particular case study, we specifically focused on L5 intratelencephalic (IT) neurons, and we categorized each cell by its highest module activity score. Notably, this approach led to the identification of three gene co-expression modules, with the highest scores for each module aligned with cells located in the inner, middle, and outer sections of the layer, each with different neighboring cells. For simplicity, we chose to compare and apply the rest of the analysis to the L5 IT neurons situated at the inner and outer sections of the layer (Figure 3B).

Following the steps described in the previous case study, we first investigate how different localization of neurons might generally affect gene expression. Despite the initial observation that the two neuron groups showed differences in gene expression, the global analysis of the cosine distances was not sufficient to elucidate differences among neuron populations (Figure 3C), showing a minimal variance in within and across groups distance distributions (median cosine distances of 0.11 and 0.12, respectively). However, UMAP projections of scRNA-seq data successfully separated the two groups, suggesting spatial gene expression differences when the 2,000 most variable genes are visualized in a two-dimensional framework (Figure S10). Indeed, in a differential expression analysis between the neuron groups, we identified 917 genes whose expression levels changed based on their location within the neuron layer, in line with the gene module analysis (Figure 3D; Table S1). Remarkably, and in agreement with the original CellTrek results, neurons situated in the inner region of the layer showed significant activity for COL27A1 and COL6A1 (logFC of 10.69 and 4.7, respectively), whereas neurons in the outer section were active in WHRN and ENDOU (logFC of 11.4 and 10.98).

To investigate the interaction dynamics between L5 neurons located at different borders of the layer (proximal to L4 or L6 layers) and the other defined cell types, we applied CellPhoneDB to this dataset. Since this is murine brain tissue, we first identified the human orthologues to the mouse genes using BioMart^30^. Next, we ran CellPhoneDB to analyze interactions between the two spatially defined neuron groups and the remaining cells in the dataset. In this case, the analysis predicted a notably high number of ligand-receptor interactions, exceeding those observed in the breast cancer dataset, with 9,983 significant interactions. Notably, L4 neurons were predicted to have the highest interaction rates with both inner and outer L5 IT neurons (Figure 3E, solid line) and, as in the breast cancer samples, we found a strong correlation comparing both neuron groups and the rest of cell types (ρ=0.82).

As in the previous case study, we assessed whether the cell types predicted to interact are located in the proximal regions of the tissue slide. To adapt the CellTrek spatial localization results for spot-level analysis, we converted the cell locations to their nearest spatial spots using Euclidean distance, allowing the accurate assignment of each cell to its closest spot. Next, we quantified the colocalization between the two neuron groups and the other cell types in the datasets. As expected, L5 IT neurons situated in the outer section predominantly colocalize with neurons from the previous L4 layer. Conversely, L5 IT neurons located in the inner section showed higher colocalization scores with cells from the subsequent L6 IT layer (Figure 3E, dotted line). This analysis reveals that neurons within the same layer can have different neighboring cells, potentially contributing to the observed variations in gene expression patterns. However, we again observed no correlation between the colocalization of neurons with other cell types and the number of predicted ligand-receptor interactions (Figure 3F). Notably, as observed in the breast cancer dataset, we identified several cases in which cells did not share spatial location yet still exhibited a high number of predicted ligand-receptor interactions. While predictions accurately reflected higher counts of LR interactions between outer layer neurons and L4 neurons, the predicted interaction counts with inner layer neurons were unexpectedly high, despite the lack of colocalization (467 and 402, respectively). This discrepancy was particularly evident in the case of interactions involving L6 IT neurons. Indeed, CellPhoneDB predictions suggested higher interaction rates with neurons in the outer section (288 and 379, respectively) than with L5 neurons colocalizing in the inner part of the layer. Additionally, there were some cases where L5 IT neurons showed no colocalization, yet a high number of LR interactions was predicted, such as with Vip or L2/L3 IT neurons (794 and 755, respectively). This result highlights a misalignment between the predicted interactions and the observed spatial colocalization patterns, despite their capacity to accurately identify expected interactions (Figure 3E, F).

These findings were further validated using the LIANA consensus approach, which confirmed similar interaction trends; L5 IT neurons in the outer part were generally predicted to have more interactions (Figure S11). However, LIANA predicted a much lower total number of LR interactions with our neurons of interest (1,907 predicted interactions with L5 IT neurons), suggesting more stringent criteria to select significant interactions, which likely yields fewer false positives. To confirm that the observed interaction patterns were not a result of the higher neuron counts in the outer part of the layer, we also repeated the subsampling analysis, ensuring the robustness of our conclusions (Figure S12).

### Case study 3. Explore fibroblasts in different tissue regions

We performed a last exercise on the Human Heart Atlas^31^, a comprehensive dataset presenting more than 700K cells and 42 different sample slides, containing 11 slides from the sinoatrial node (SAN) and 8 from the atrioventricular node (AVN). We focused on the SAN and AVN slides due to their distinct histological annotations within the same slide provided in the original publication, indicating specific cellular niches within each slide, and we assessed whether cells positioned in different histological regions of the same slide exhibit distinct communication patterns due to their unique microenvironments.

We first mapped cells to their respective spatial slides (Figure 4A and S13). The authors provided results from deconvolution using cell2location^32^, which showed partial agreement with our cell mapping results. The observed discrepancies could be attributed to their use of all available cells from a region to define a reference cell type prior to deconvolution, whereas we only used patient-slide-matched cells for mapping. Notably, we were unable to map P cells into the histologically annotated nodes using any methodology, with Tangram yielding the most different results among the mapping techniques (Figure S14). Overall, we annotated eleven different cell types in both spatial slides, including adipocytes, atrial and ventricular cardiomyocytes, lymphatic endothelial and endothelial cells, fibroblasts, lymphoid cells, mast cells, mural cells, myeloid cells, and neural cells. Examination of cell abundances by slide (Figure S15) showed similar cell proportions across slides, except for ventricular cardiomyocytes exclusive to the AVN slide and a substantial difference in lymphoid cell presence, with the SAN slide showing higher counts in lymphoid cell populations. Predicted cell type abundances across different regions revealed significant compositional variations based on histological region (Figure S16). For instance, fibroblasts were predominantly found in the node and the myocardium for both slides. Moreover, the node region on the SAN slide showed a more diverse cell composition compared to the node region in the AVN slide, which was largely dominated by fibroblasts.

**Figure 4.**
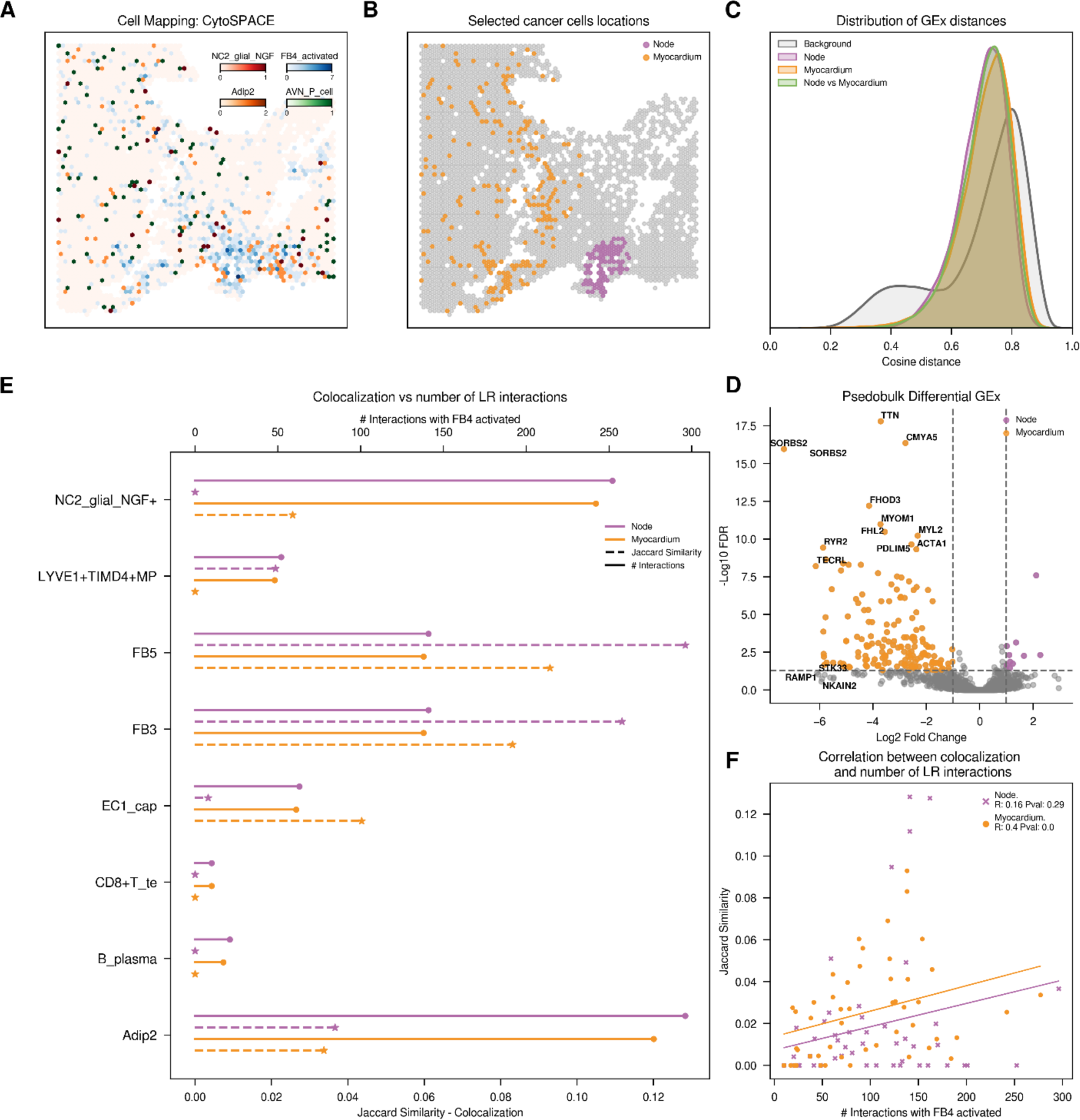
Exploration of fibroblast (FB4 activated) from ANV slide. **A.** Distribution of main cell types across the tissue slide. For visualization purposes, we normalized the CytoSPACE inferred abundances of predominant cell types, with each spatial spot representing the cell type of highest abundance. **B.** Spatial representation of fibroblasts (FB4 activated) according to their assigned location **C.** Distribution of cosine distances between gene expression (GEx) profiles. We analyze the cosine distances between gene expression patterns of fibroblasts located within the same or different tissue histological regions. **D.** Volcano plot with Differential Gene Expression results, highlighting significant upregulated genes in fibroblast of each histological region. **E.** Comparison of colocalization and CellPhoneDB cell-cell communication results. Plain line (top axis) represents the count of significant ligand-receptor interactions between each of the defined fibroblasts and the y-axis cells. Dotted line (bottom axis) indicates colocalization, measured by the Jaccard similarity index based on the presence or absence of cells within each spot. **F**. Spearman correlation between colocalization and ligand-receptor counts Color code: Violet fibroblast in the node and orange fibroblasts in the myocardium.

We identified Fibroblast 4 Activated (FB4) cells in both the node and in adjacent histological areas like the myocardium and epicardium, allowing the exploration of their communication and colocalization patterns across these regions (Figure 4B). Analysis of cosine distances between gene expression profiles of FB4 cells showed no differences within or between different histological locations (Figure 4C). Furthermore, the medians of these distributions indicated a lack of similarity among the FB4 cells within the same region (AVN: 0.72 and SAN: 0.75), suggesting heterogeneity in their gene expression. As in the breast cancer dataset, UMAP projections (Figure S17) of scRNA-seq data supported the lack of gene expression difference between spatial groups, where cells failed to cluster by histological location. In addition, the results from differential gene expression analysis were ambiguous, with 195 genes showing location-dependent differential expression in the AVN sample and not a single gene differentially expressed in the SAN sample (Figure 4D; Table S1).

Finally, we explored the relationship between colocalization and predicted intercellular communication (Figure 4E and 4F). A substantial number of significant LR interactions were identified for both slides (AVN: 10,515 and SAN: 8,077 interactions involving FB4 and other cells in the dataset). Again, the correlation between the most interactive cell types across the two histological groups compared was strong (AVN: 0.96 and SAN 0.94), while the correlation between colocalization was weaker (AVN: 0.48 and SAN: 0.36). CellPhoneDB found very low counts of LR interactions between e.g. T cells CD8 and plasma B cells, which did not colocalize with any defined histological regions in the AVN slide. However, endothelial cells (EC1), despite having a poor colocalization with FB4 in the node, were predicted to interact with FB4 at similar frequencies than EC1 present in the myocardium, where colocalization with fibroblast did occur (63 and 61, respectively). Similar results were observed with tissue-resident macrophages (LYVE1^+^IGF1^+^ MPs), while in this case the colocalization was with the node and not the myocardium and the predicted number of LR interactions was practically the same (59 and 58 LR interactions, respectively). In the SAN slide, we also identified a low number of LR interactions between fibroblasts and B or mast cells, which are not found to colocalize. However, once again, we observed some discrepancies in, for instance, monocytes (CD4^+^Mo) or innate lymphoid cells (ILC) which, despite the lack of colocalization with FB4 in one of the defined regions, exhibited an identical number of predicted interactions. The robustness of these findings was further validated through subsampling analysis (Figure S18) and also using LIANA to predict cell-cell communication (Figure S19).

## Concluding remarks

Understanding the communication between cells and their microenvironment is fundamental to unravel the cellular complexity of tissues and, ultimately, organ function. Currently, RNA sequencing is the only technology scalable enough to provide detailed molecular information at single-cell resolution and that can be applied to tissue slides of significant size. Accordingly, to date, most of the efforts to study the molecular bases of cell-cell communication rely on single-cell transcriptional data. In this article, we have implemented a computational strategy that integrates cellular spatial organization with single-cell transcriptomics to systematically investigate how different tissue microenvironments affect the cellular gene expression and intercellular interactions. We have applied our approach to many samples in three different tissues (i.e. breast, brain and heart), identifying multiple cell coalitions embedded in niches with different cellular compositions. Our analyses reveal that the global transcriptional profile of a cell type does not significantly change depending on the cellular composition of its microenvironment, with neither profile similarity distributions nor UMAP representations being able to distinguish between cells in different niches. In addition, when we compared the gene expression between the same cell types in different locations, we find differentially expressed genes in less than half of the samples analyzed. Together, these observations severely limit the ability of current cell-cell communication methods, based on the detection of ligand-receptor expression, to account for subtle differences in gene expression influenced by different microenvironments. Indeed, when comparing the results of such methods, we find almost perfect correlations between the cell-cell contacts predicted in the different niches, where the cell composition is quite different. Moreover, we see how these models are not capable of recapitulating spatial cell distribution and distinguishing diverse communication dynamics.

Mapping single cells onto distinct tissue locations remains an inference process, where gene expression profile matching is used to assign each cell to a specific spot. State-of-the-art use global optimization methods to align single-cell and spatial transcriptomics, which is already improving their accuracy^19^. However, in the absence of a clear set of differentially expressed genes, the allocation of cells to specific locations or niches can be challenging, resulting in less-organized tissue areas. An increased availability of spatially resolved transcriptomics datasets are prompting the development of novel strategies to infer cell-cell communication which readily incorporate information on the cells location, providing a better inference of ligand-receptor pairs^7,15,33^. It thus seems clear that next-generation methodologies will successfully identify cellular locations. However, the lack of detectable differences in gene expression between cells of the same type embedded in different microenvironments questions the appropriateness of using single-cell 10x sequencing data, together with generic lists of receptor-ligand pairs, as only proxy to predict cell-cell communication. We believe that current cell-cell contact inference approaches are, indeed, useful to suggest lists of proteins mediating such interactions, but they are not sensitive enough to capture context-specific differences. And, to develop approaches that identify the molecular bases of *bona fide* functional interactions between cells, we need to acquire a quantitative understanding of the factors governing cell-cell communication.

## Datasets and Methods

### Datasets

#### Breast Cancer Atlas

The original Breast Cancer Atlas publication generated scRNA-seq data from 26 primary tumors, including 11 ER^+^, 5 HER2^+^, and 10 TNBC tumors^20^. A total of 130,246 single cells passed the quality controls. We pre-processed the dataset using the standard Scanpy pipeline and followed the authors’ recommendations ^34^. In brief, we normalized gene expression counts, identified highly variable genes (n = 2,000), performed principal component analysis (n = 100), neighbor identification, and Leiden clustering. We annotated the clusters using the cellular subtypes described in the original publication. To integrate this data with spatial information, we selected cells from patients for which we had spatial information (19,300 cells). The dataset contained spatial slides on two ER^+^ tumors, lobular and ductal, respectively (CID4535 and CID4290), and two TNBC ductal tumors (CID44971 and CID4465). Again, we pre-processed the spatial data using the standard Scanpy pipeline for Visium datasets. Briefly, we filtered spots based on total counts (5,000 < counts < 35,000) and percentage of mitochondrial counts (20%). In addition, we filtered genes that were expressed in less than 10 spots. Finally, we normalized using Scanpy default parameters and detected highly variable genes (2,000). As for scRNA-seq data, we continued with the PCA, neighbor, and Leiden clustering. Clusters and pathological annotation of each spot were represented on the H&E images.

#### Mouse Brain Cortex

We downloaded scRNA-seq data representing adult mouse cortical cells from the Allen Brain Atlas^27^ (https://www.10xgenomics.com/datasets/mouse-brain-serial-section-1-sagittal-anterior-1-standard-1-1-0), which was generated utilizing the SMART-seq2 protocol. Additionally, we sourced a spatial transcriptomics (Visium v1) slide from the anterior section of the sagittal mouse brain. We downloaded both single-cell and spatial transcriptomics datasets as pre-processed Seurat objects from the CellTrek GitHub^3,4^, and we obtained the cell type annotations directly from the data.

#### Human Heart Atlas

We downloaded processed sn/scRNA-seq data from the Human Heart Altas^31^ (https://www.heartcellatlas.org/). In brief, they collected 704,296 cells and nuclei from eight anatomical cardiac regions, including the cardiac conduction system. They profiled 25 healthy hearts and generated 42 Visium slides, most from the sinoatrial (SAN) and atrioventricular (AVN) nodes. sc/snRNA-seq and spatial transcriptomics data were pre-processed by the authors, so cell type annotations or histological annotations were directly obtained from the Scanpy AnnData object. Additionally, in the spatial AnnData object, the authors provide the predicted abundance of each defined cell state in each spot, estimated using cell2location^32^. For our exercise, we selected scRNA-seq data from subjects AH1 and A61 (13,019 and 20,995 cells, respectively) and corresponding SAN and AVN spatial slides (HCAHeartST10659160 and HCAHeartST11290662).

### Deconvolution and cell mapping: scRNA-seq and ST integration

To spatially map cell types defined by scRNA-seq data and delineate regions with diverse cellular compositions, we combined deconvolution and cell mapping methodologies to integrate scRNA-seq and ST measurements. Specifically, we utilized Cell2location^32^ for deconvolution, enabling the estimation of the expected abundances of various cell types across the spots. In parallel, we employed three cell mapping techniques (CellTrek^5^, CytoSPACE^19^, and Tangram^21^) to precisely assign individual cells to spatial locations.

#### Cell2location

Following the protocol defined in the Cell2location GitHub repository ^32^ https://github.com/BayraktarLab/cell2location, we first generated reference signatures of cell types using scRNA-seq data from each specific patient or sample using a negative binomial regression model provided in the Cell2location package. For every spatial slide, we built patient-specific reference signatures using raw RNA counts. Next, the inferred reference signatures were employed to estimate the abundance of each cell type within each Visium spot by decomposing the RNA counts of the spots. The regression model was trained using parameters: max_epochs = 250, batch_size = 2,500. For spatial mapping, we estimated five cells per location, which is standard for Visium datasets, set detection_alpha at 20, and increased max_epochs to 30,000.

#### CellTrek

CellTrek allows the mapping of single cells to their spatial coordinates by aligning scRNA-seq with ST data^5^. This algorithm utilizes Spatial Seurat for coembedding scRNA-seq and ST data^35^, followed by a random forest model to predict spatial coordinates. Following the procedure detailed in the documentation available on the CellTrek GitHub repository (https://github.com/navinlabcode/CellTrek), we ran CellTrek with default settings using log2-normalized data. To ensure that each cell was uniquely assigned to a single location without cell duplication, we modified the top_spot parameter to 1. For colocalization analysis and comparison with other mapping methodologies, we selected the closest ST spot for each cell via Euclidean distance.

#### CytoSPACE

CytoSPACE is a method for assigning single-cell transcriptomes to spatial transcriptomics spots via shortest augmenting path optimization^19^. Distinct from CellTrek, CytoSPACE estimates cell type fractions within the ST data. This method utilizes Seurat to assess the overall cell type composition across all the spots, but it is compatible with the output from other deconvolution methods, including Cell2location. Additionally, CytoSPACE estimates the number of cells per ST spot, enabling the sampling of scRNA-seq data to match the estimated cell counts per cell type. Following the authors’ recommendations: https://github.com/digitalcytometry/cytospace, we ran CytoSPACE using raw RNA counts and default parameters.

#### Tangram

Tangram is an integration method to combine spatial transcriptomics and single-cell RNA-seq data^21^. This technique utilizes non-convex optimization alongside deep learning to learn a spatial alignment for single-cell data, enabling the assignment of cells to spatial locations in histological sections. We followed the procedure outlined in the Tangram GitHub repository (https://github.com/broadinstitute/Tangram). We started by selecting common genes between spatial and scRNA-seq log-normalized data. Then, we trained the model to map annotated cells to the spatial spots (num_epochs = 500, density_prior = “rna_count_based”). Tangram generates a probability prediction of cell-spot association^36^. We refined this output by selecting only the most probable spot assignments for each cell.

### Downstream spatial analysis

To investigate the impact of cellular niche and cellular composition on cell type-specific gene expression, we performed several downstream analyses focusing on the spatial distribution of selected cells. In these analyses, selected cells were categorized into distinct groups based on their spatial positioning, reconstructed by previously described methodologies. Group division criteria varied depending on the analyzed system, as detailed in each of the case studies. We explored differences between spatial groups at different resolutions.

#### Cosine distances between GEx profiles

To assess overall similarities in gene expression profiles in relation to cell spatial locations, we computed pairwise cosine distances between normalized gene expression vectors. Distance calculations were performed for cells both within the same spatially defined group and across different groups. The resulting cosine distance distributions were then compared using the Kolmogorov-Smirnov test.

#### Differential Gene Expression Analysis

To identify differentially expressed genes (DEGs) within the specified cell type across defined spatial groups, we employed two complementary approaches. Initially, DEGs were identified using the rank_genes_groups() function from Scanpy, which utilizes the non-parametric Wilcoxon rank-sum test for differential expression testing. P-values obtained from this analysis were corrected for multiple comparisons using the Benjamini-Hochberg correction. Genes were considered significantly differentially expressed at an absolute fold change greater than 1.5 and an adjusted p-value below 0.05.

Subsequently, to address potential challenges associated with single-cell data^37^, such as zero inflation and high cell-to-cell variability, we adopted a pseudobulk approach. By aggregating cell expression data into ‘pseudo-replicates’, we applied a Python implementation of the DESeq2 method (pydeseq2), designed for bulk expression samples^38,39^. This method mitigates p-value inflation and augments robustness. For this analysis, pseudoreplicates were constructed using raw gene expression (n=3 for condition), and differential expression was assessed using the Wald test. P-values were again adjusted using the Benjamini-Hochberg method, and we applied the same criteria for significant differential expression (absolute log fold change > 1.5 and adjusted p-value < 0.05).

To relate identified DEGs with their biological functions, we conducted enrichment analysis at the GO Biological Process 2023, GO Molecular Function 2023^40^, and Reactome 2022^41^ through the Enrichr API^42^. We selected only terms with adjusted p-values <0.05. We employed REVIGO to summarize the GO terms^43^, thereby enhancing our comprehension of the different mechanisms being altered.

#### Colocalization analysis

To investigate the spatial colocalization of different cell types based on mapping results, we adapted a previously defined strategy to quantify their spatial overlap^19^. First, we constructed a spot-presence binary matrix, where each spot was scored as 1 if at least one cell of a particular cell type was present and 0 otherwise. We then used the columns of this matrix, representing the spot profiles for each cell type, to calculate pairwise Jaccard similarities. This similarity metric assessed the spot overlap between two cell types, with higher Jaccard values referring to a greater tendency for colocalization of these cells in the space.

By applying this colocalization metric, we were able to quantitatively characterize unique microenvironments for the previously defined cell groups, enabling the examination of how colocalization varies across different spatial regions. Moreover, this approach also offers a ground truth for cell-cell communication studies. Given that intercellular interactions are predominantly spatially constrained, cells exhibiting higher colocalization have higher chances of interaction.

### Intercellular communication

To analyze intercellular communication, we prepared log-normalized counts and metadata relating cell barcodes to previously defined cell labels as our input data. As previously mentioned, we identified a cell type of interest located in two distinct tissue regions, which were accordingly labeled differently. This differentiation allowed us to investigate potential variations in communication patterns within the same cell type across different spatial regions, aiming to correlate these differences with the colocalization analysis.

#### CellPhoneDB

CellPhoneDB is one of the core tools for studying cell-cell communication^8^. It is a statistically based tool that uses permutation to calculate the significance of a predicted interacting pair. CellPhoneDB is based on the coordinate expression of ligands and receptors to infer cell interactions using expression mean as a scoring function. For our study, we utilized the Python implementation of CellPhoneDB v.4, setting the significant threshold at 0.05 to identify statistically significant interactions.

#### LIANA

To extend our analysis beyond CellPhoneDB and systematically assess cell-cell communication across various computational methods, we utilized LIANA, an integrated framework that facilitates the exploration of interactions inferred by seven distinct computational tools^26^. LIANA addresses the output heterogeneity among tools by incorporating a consensus prediction, which provides more reliable results. We run LIANA using default parameters, with resource_name = ‘consensus’, min_cells = 5, expr_prop = 0.1, and n_perms = 100.

#### Subsampling analysis

To accurately compare and control for the potential impact of cell counts on our analysis of cell-cell communication differences among the same cell type across different tissue locations, we implemented a subsampling strategy. This approach aimed to normalize the number of cells associated with each spatial label, thereby preventing any bias due to variable cell counts. First, we identified the group with the lowest cell count and then subsampled the other group to match this number. To mitigate any sampling biases introduced by this procedure, we built 10 unique subsampled datasets. Following this, we proceeded with the cell-cell communication analysis on each of these datasets, ensuring that our predictions were not affected by the discrepancies in cell numbers.

## Code availability

All the code necessary to replicate the analyses of the three different systems is available in our GitLab repository (https://gitlabsbnb.irbbarcelona.org/epareja/cell-cell-communication.git). The repository contains Jupyter Notebooks for user convenience, along with detailed explanations of the repository content, required data, and all results showcased in the manuscript.

## Acknowledgments

P.A. acknowledges the support of the Generalitat de Catalunya (RIS3CAT Emergents CECH: 001-P-001682 and VEIS: 001-P-001647; and 2021 SGR 00876), the Spanish Ministerio de Ciencia, Innovación y Universidades (PID2020-119535RB-I00; PID2023-152296OB-I00), the Instituto de Salud Carlos III (IMPaCT-Data), and the European Commission (CLARITY: 101137201). E.P-L. is a recipient of an FI fellowship (2022 FI_B_00767), which is co-financed by the European Union through the European Social Fund Plus (ESF+). We also acknowledge institutional funding from the Spanish Ministry of Science and Innovation through the Centres of Excellence Severo Ochoa Award, and from the CERCA Programme / Generalitat de Catalunya.

## Notes

### Competing Interest Statement

The authors have declared no competing interest.

