## Supplementary Figures for "Systematic assessment of microenvironment-dependent transcriptional patterns and intercellular communication"

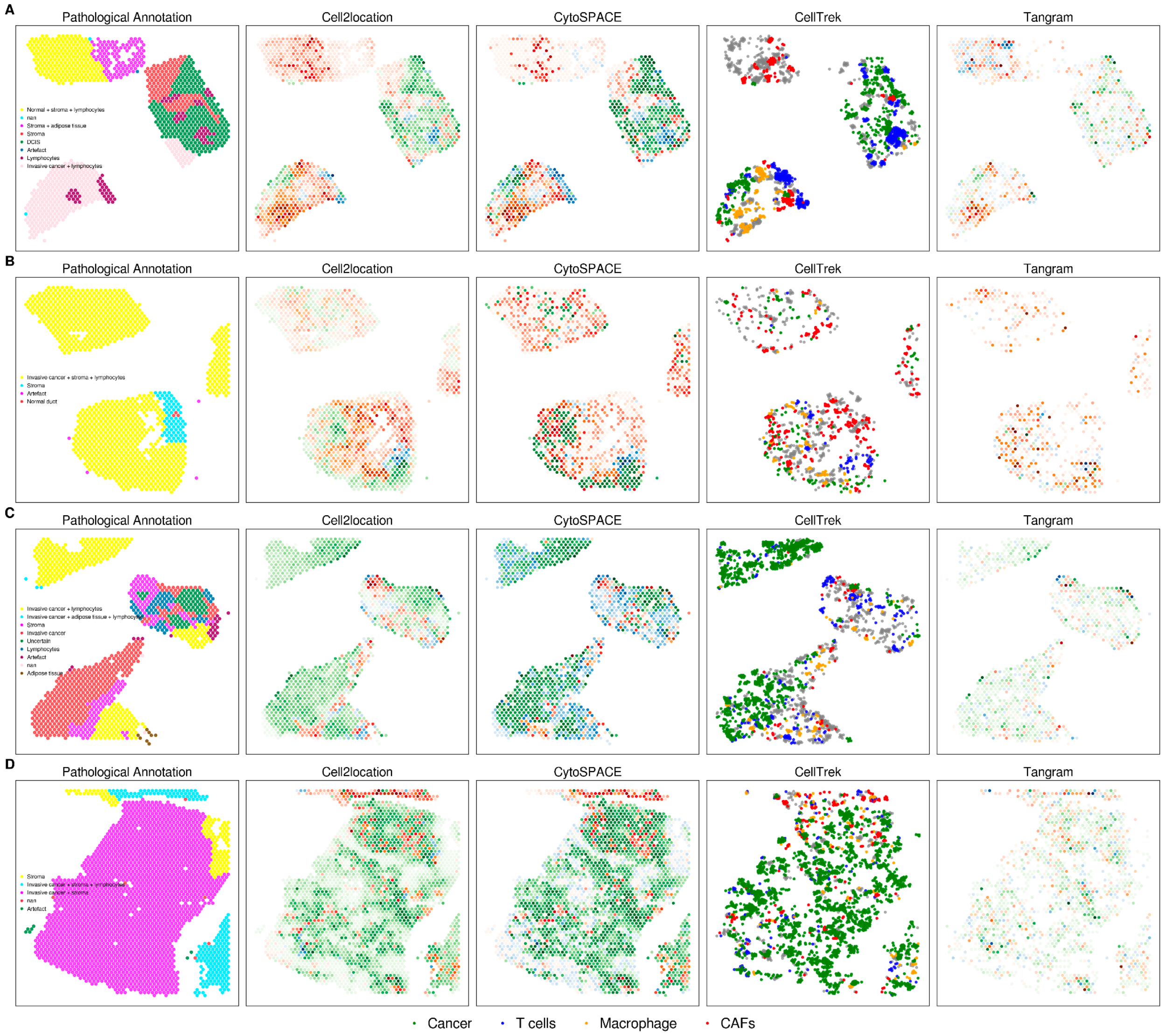


**Figure S1. Methodological comparison for disentangling single-cell spatial resolution for breast cancer dataset**. We applied four methodologies - Cell2location, CytoSPACE, CellTrek and Tangram - to determine cell positioning across different patients (identified as CID44971, CID4465, CID4535, CID4290, and labeled A, B, C, D respectively). Pathological annotations from the original publication are included for comparative analysis (Wu et al., 2021), showing notable concordance between the methodologies and the annotations. The figure illustrates the normalized abundance of predominant cell types, with each spatial spot representing the cell type of highest abundance for clear visualization. Color coding: green cancer cells, blue T cells, red macrophages and orange Cancer Associated Fibroblasts (CAFs).

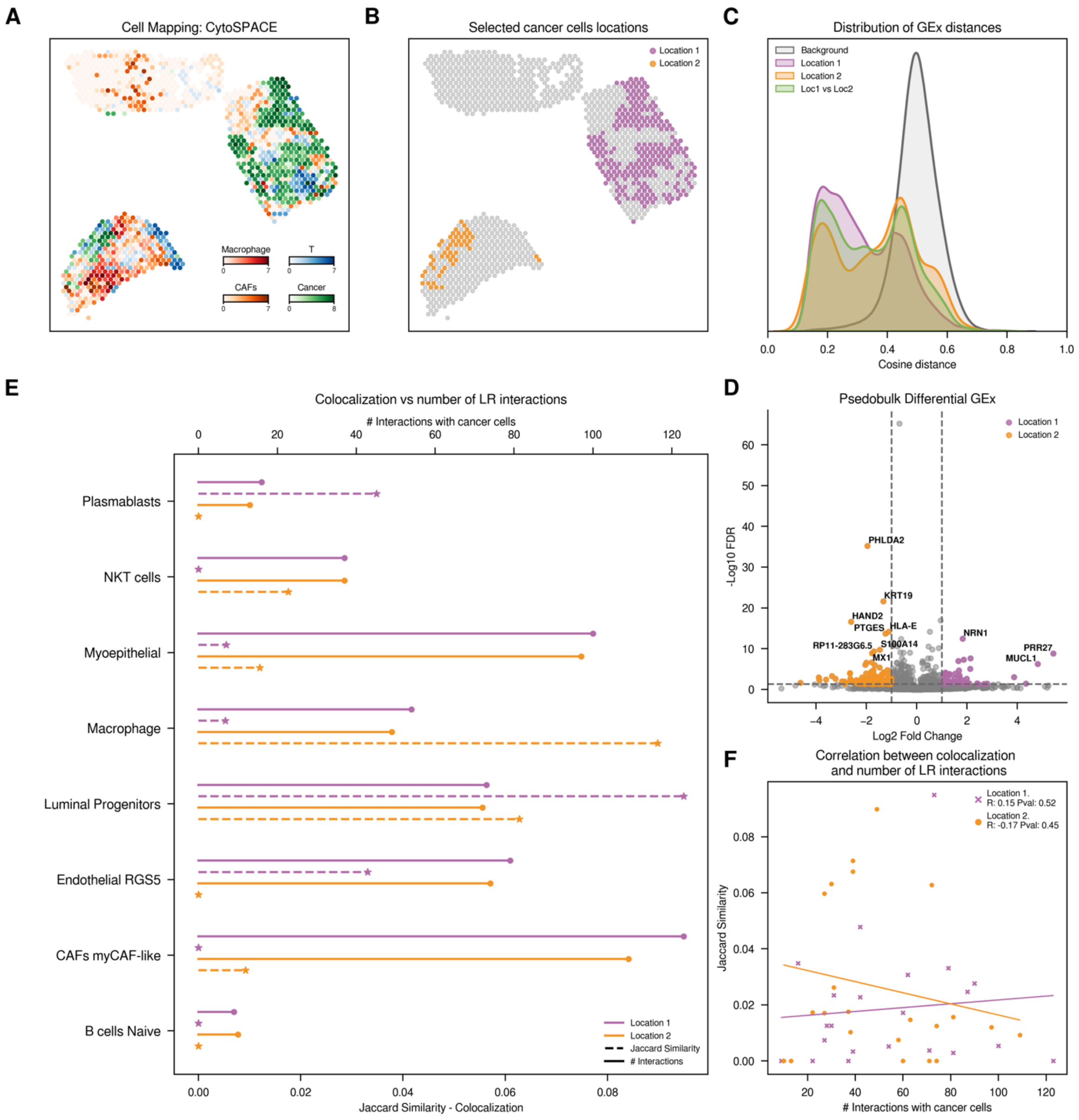


**Figure S2. Exploration of cancer cells from CID44971 patient. A.** Distribution of main cell types across the tissue slide. For visualization purposes, we normalized the CytoSPACE inferred abundances of predominant cell types, with each spatial spot representing the cell type of highest abundance. **B.** Spatial representation of cancer cells according to their assigned location. **C.** Distribution of cosine distances between gene expression (GEx) profiles. We analyze the cosine distances between gene expression patterns of cancer cells located within the same or different tissue regions. **D.** Volcano plot with Differential Gene Expression results, highlighting significant upregulated genes in cancer cells of each region. **E.** Comparison of colocalization and CellPhoneDB cell-cell communication results. Plain line (top axis) represents the count of significant ligand-receptor interactions between each of the defined cancer cells and the y-axis cells. Dotted line (bottom axis) indicates colocalization, measured by the Jaccard similarity index based on the presence or absence of cells within each spot. **F**. Spearman correlation between colocalization and ligand-receptor interaction counts. Color code: Violet cancer cells in location 1, orange cancer cells in location 2.

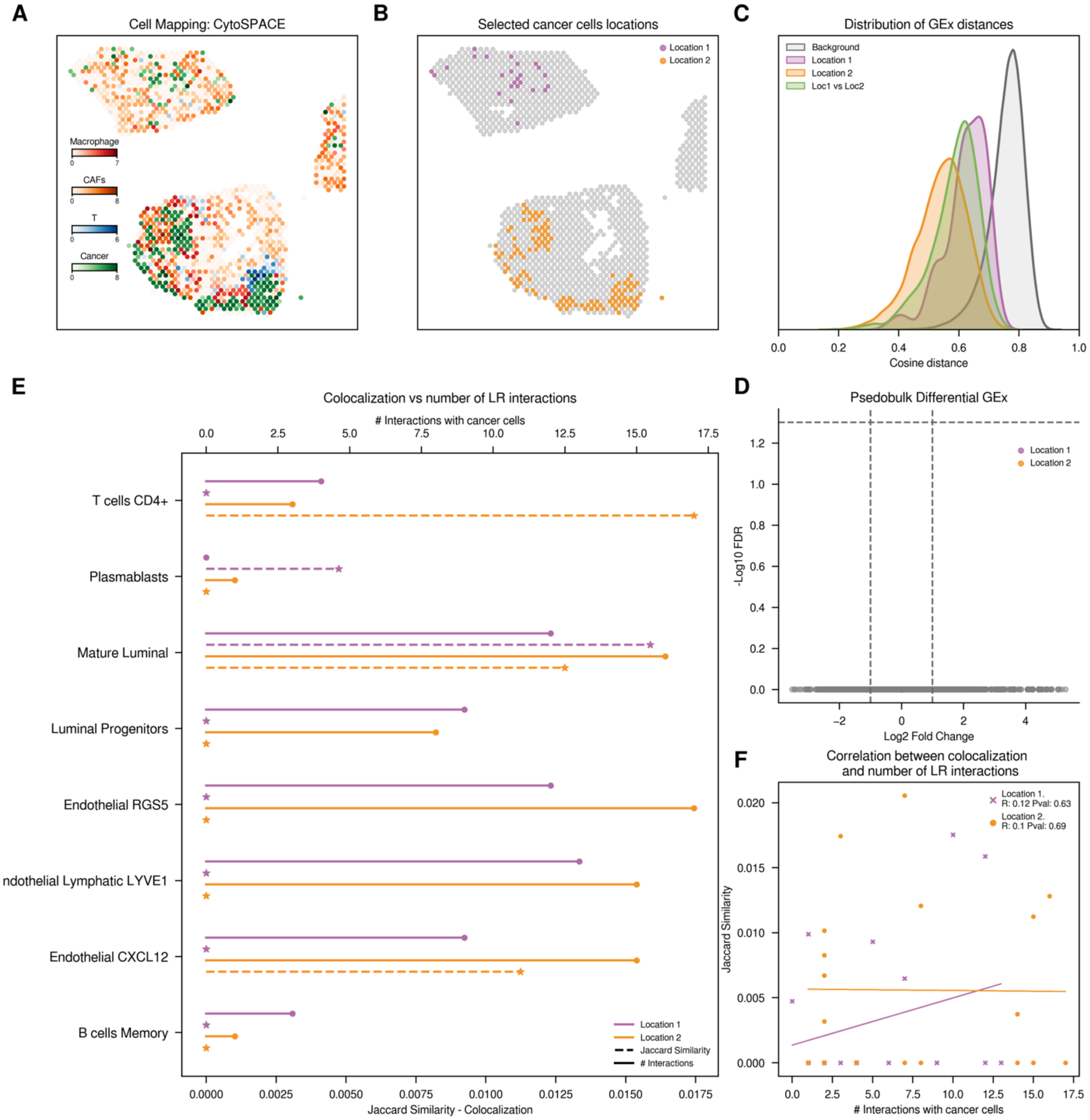


**Figure S3. Exploration of cancer cells from CID4465 patient. A.** Distribution of main cell types across the tissue slide. For visualization purposes, we normalized the CytoSPACE inferred abundances of predominant cell types, with each spatial spot representing the cell type of highest abundance. **B.** Spatial representation of cancer cells according to their assigned location. **C.** Distribution of cosine distances between gene expression (GEx) profiles. We analyze the cosine distances between gene expression patterns of cancer cells located within the same or different tissue regions. **D.** Volcano plot with Differential Gene Expression results, highlighting no significant upregulated genes in cancer cells of each region. **E.** Comparison of colocalization and CellPhoneDB cell-cell communication results. Plain line (top axis) represents the count of significant ligand-receptor interactions between each of the defined cancer cells and the y-axis cells. Dotted line (bottom axis) indicates colocalization, measured by the Jaccard similarity index based on the presence or absence of cells within each spot. **F**. Spearman correlation between colocalization and ligand-receptor interaction counts. Color code: Violet cancer cells in location 1, orange cancer cells in location 2.

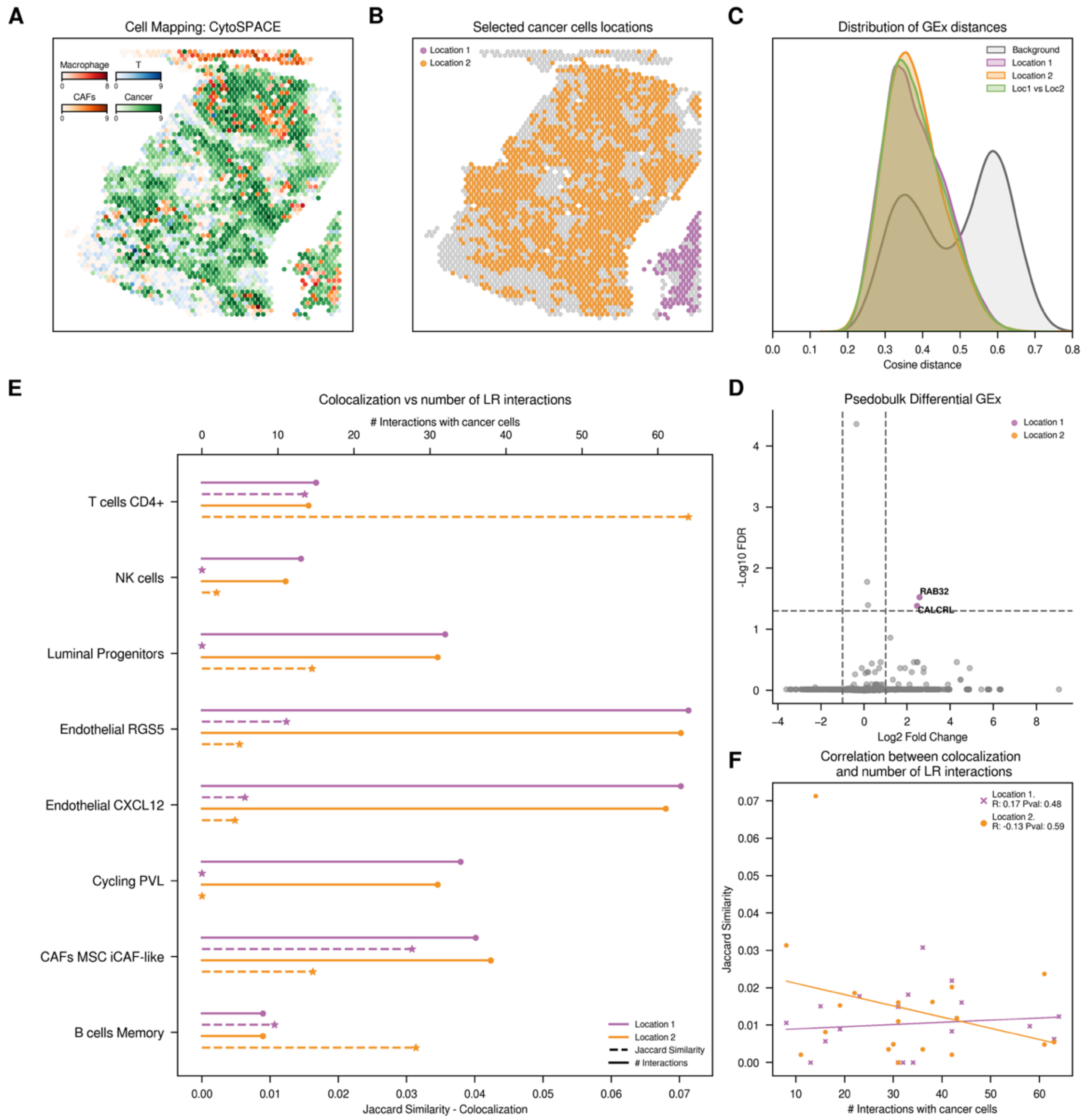


**Figure S4. Exploration of cancer cells from CID4290 patient. A.** Distribution of main cell types across the tissue slide. For visualization purposes, we normalized the CytoSPACE inferred abundances of predominant cell types, with each spatial spot representing the cell type of highest abundance. **B.** Spatial representation of cancer cells according to their assigned location. **C.** Distribution of cosine distances between gene expression (GEx) profiles. We analyze the cosine distances between gene expression patterns of cancer cells located within the same or different tissue regions. **D.** Volcano plot with Differential Gene Expression results, highlighting significant upregulated genes in cancer cells of only one region. **E.** Comparison of colocalization and CellPhoneDB cell-cell communication results. Plain line (top axis) represents the count of significant ligand-receptor interactions between each of the defined cancer cells and the y-axis cells. Dotted line (bottom axis) indicates colocalization, measured by the Jaccard similarity index based on the presence or absence of cells within each spot. **F**. Spearman correlation between colocalization and ligand-receptor interaction counts. Color code: Violet cancer cells in location 1, orange cancer cells in location 2.

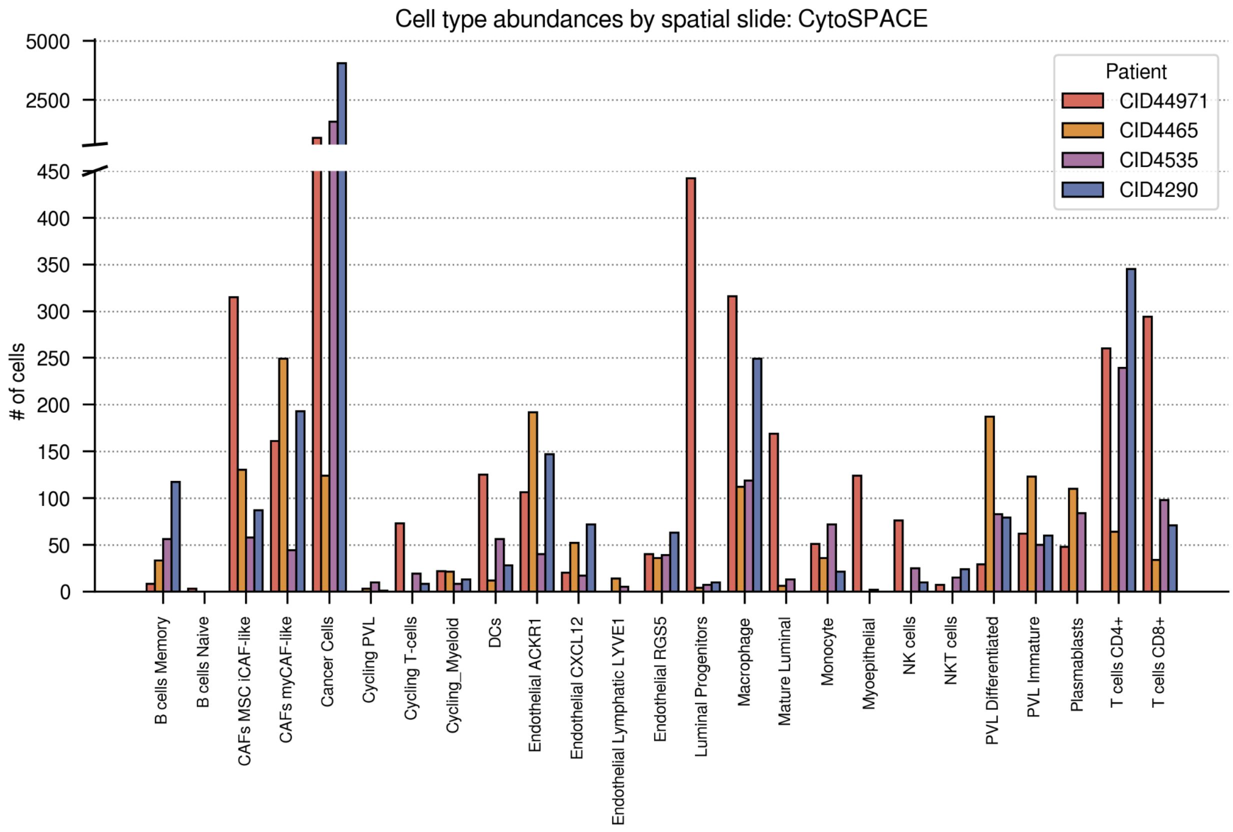


**Figure S5**. **Variability of cell type abundance across breast cancer patient tissue samples.** Bar plot showing the distribution of cell types, determined using CytoSPACE, across spatial slides from the four breast cancer patients. Abundances were calculated by summing the number of cells of each type mapped to individual spots within the CytoSPACE results. Each bar represents the count of a specific cell type within a slide, and distinct colors are assigned to each patient to facilitate comparison.

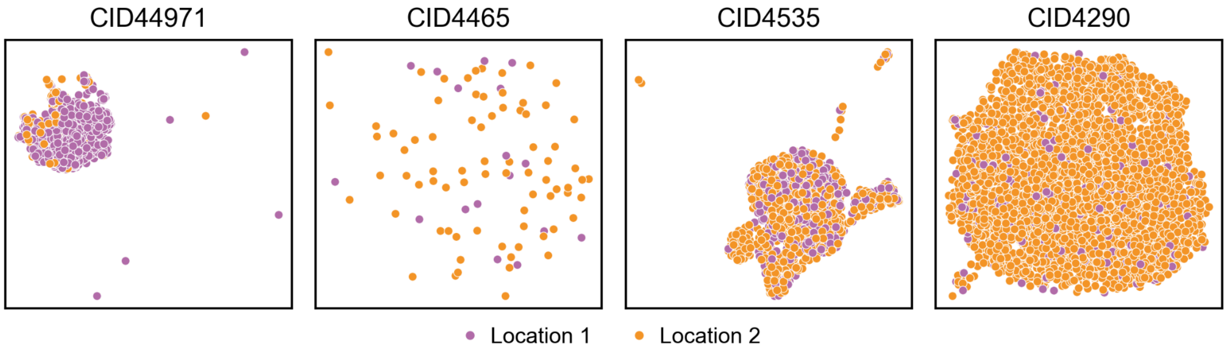


**Figure S6**. **UMAP projection of cancer cell gene expression by location.** Two-dimensional Uniform Manifold Approximation and Projection (UMAP) visualization of the gene expression profiles of cancer cells by patient, to explore the potential unsupervised separation based on the assigned location. Each point represents an individual cancer cell, colored according to its assigned location.

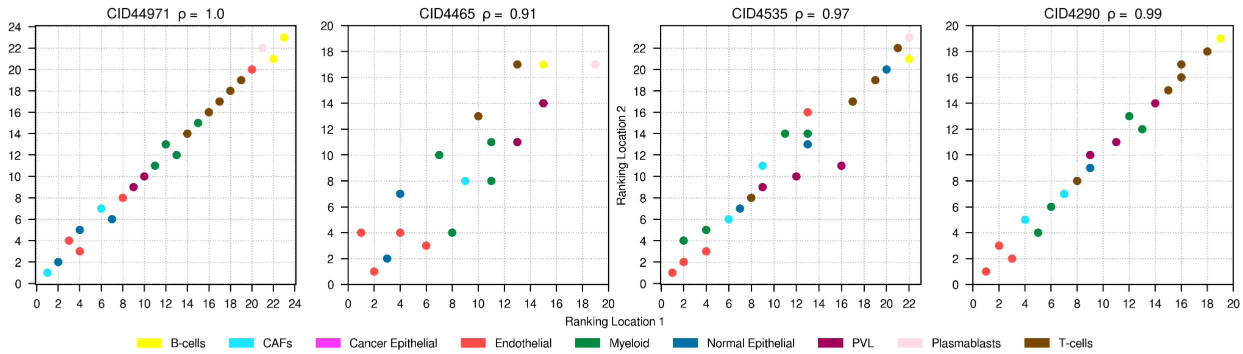


**Figure S7**. **Correlation of interaction frequency between non-cancer cells and cancer cells across different tumor regions.** Using the interactions inferred by CellPhoneDB, we analyzed the correlation in interaction frequency between non-cancer cells and cancer cells in each of the defined regions. Spearman correlation coefficients - 0.995, 0.91, 0.97, 0.99- reveal a strong correlation in interaction frequency across both groups of spatially defined cancer cells for all samples.

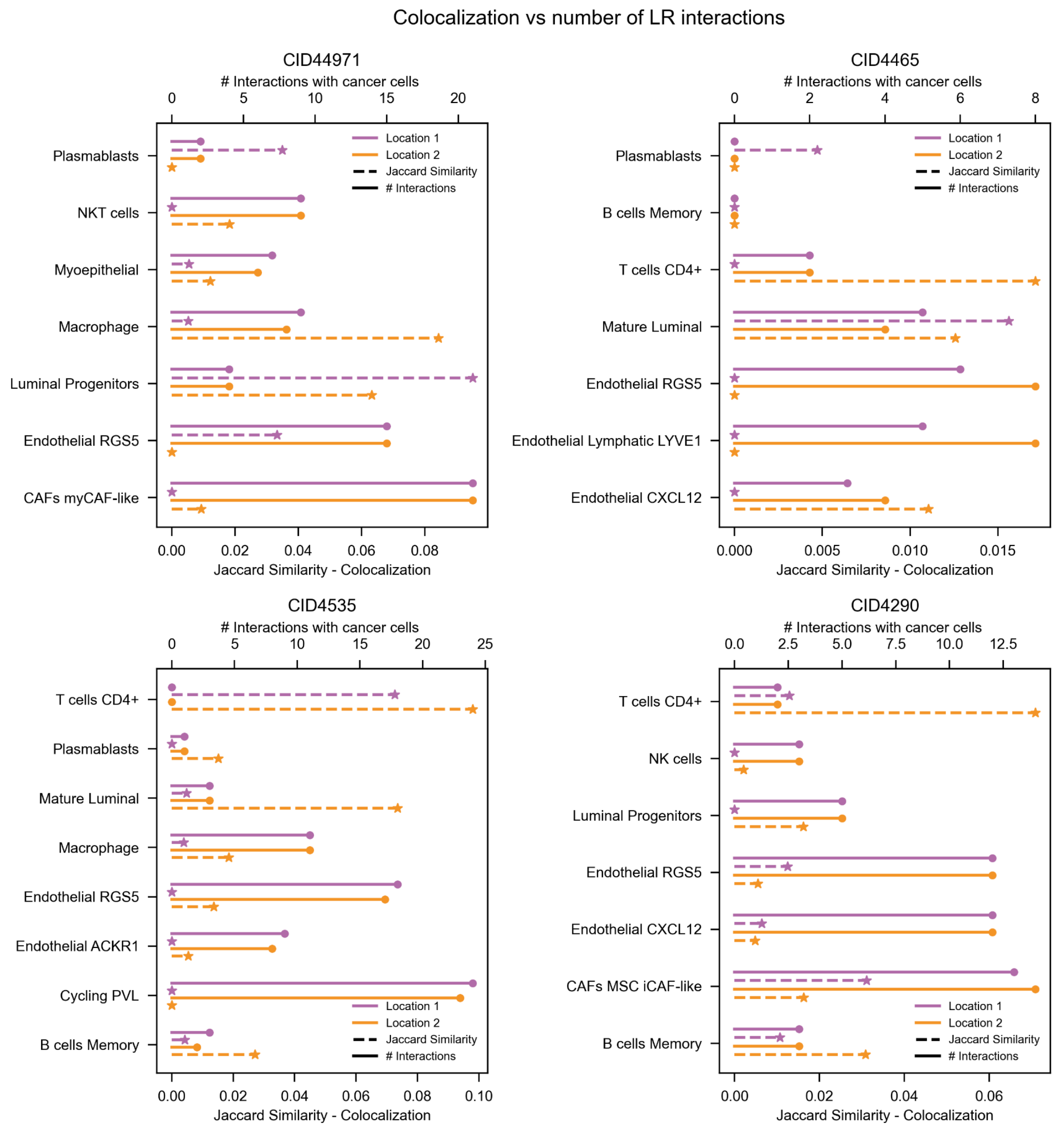


**Figure S8. Comparison of colocalization and LIANA cell-cell communication results across four breast cancer samples.** This figure presents a comparative analysis of colocalization results and the number of ligand-receptor interactions predicted between location-defined cancer cells and cells in the y-axis across four breast cancer samples. Plain line represents the number of significant ligand-receptor interactions identified by LIANA, utilizing the consensus score. Dotted line illustrates the degree of colocalization, measured as the Jaccard similarity index between the spot profiles of cancer cells and y-axis cells (based on the presence or absence of cells in each spot). For visualization purposes, we selected the extreme cases, those with the highest and lowest levels of colocalization and interaction frequencies.

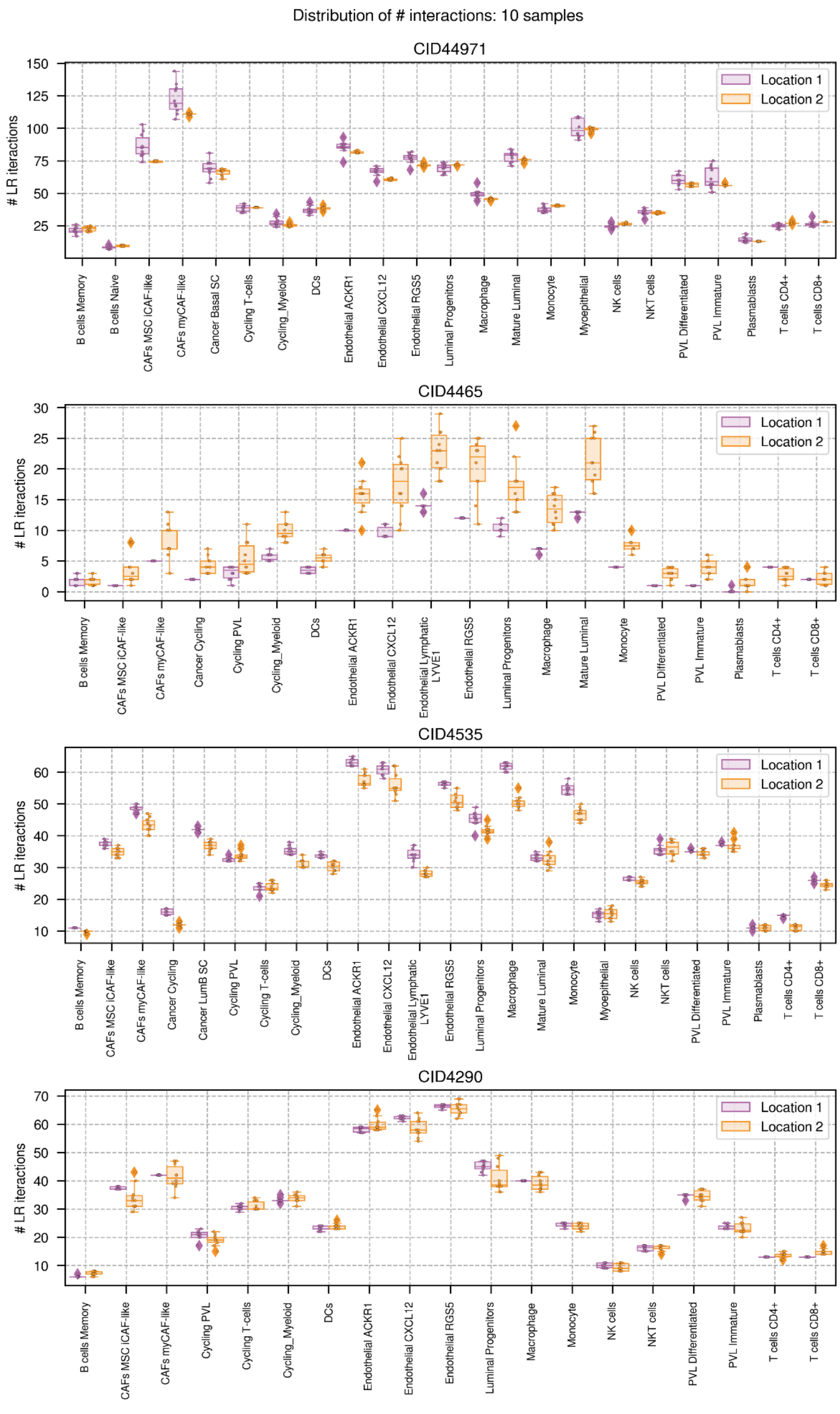


**Figure S9. Subsampling exercise in the four breast cancer samples.** This figure illustrates the minimal variability in predicted ligand-receptor interactions counts between cancer cells located in different regions and the rest of the cells in the dataset, over several subsampling iterations. To ensure a balanced analysis and mitigate biases due to varying cancer cell quantities in each region, ten unique subsampled datasets were created. Then, we predict the ligand-receptor interactions with CellPhoneDB adjusted for these quantity discrepancies.

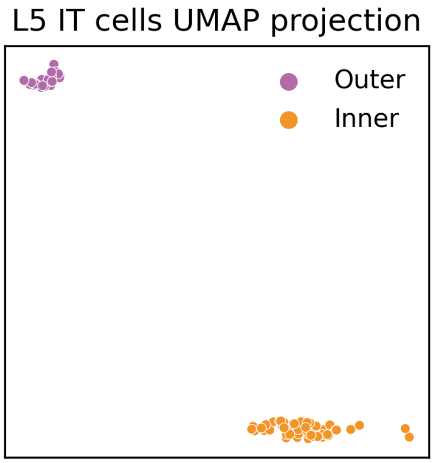


**Figure S10**. **UMAP projection of L5 IT neurons gene expression by location.** Two-dimensional Uniform Manifold Approximation and Projection (UMAP) visualization of the gene expression profiles of L5 IT neurons, to explore the potential unsupervised separation based on assigned location within the L5 layer. Each point represents an individual neuron, colored according to its assigned location. In this case, the UMAP visualization demonstrates a clear separation between the gene expression profiles of L5 IT neurons positioned in the inner versus the outer neurons.

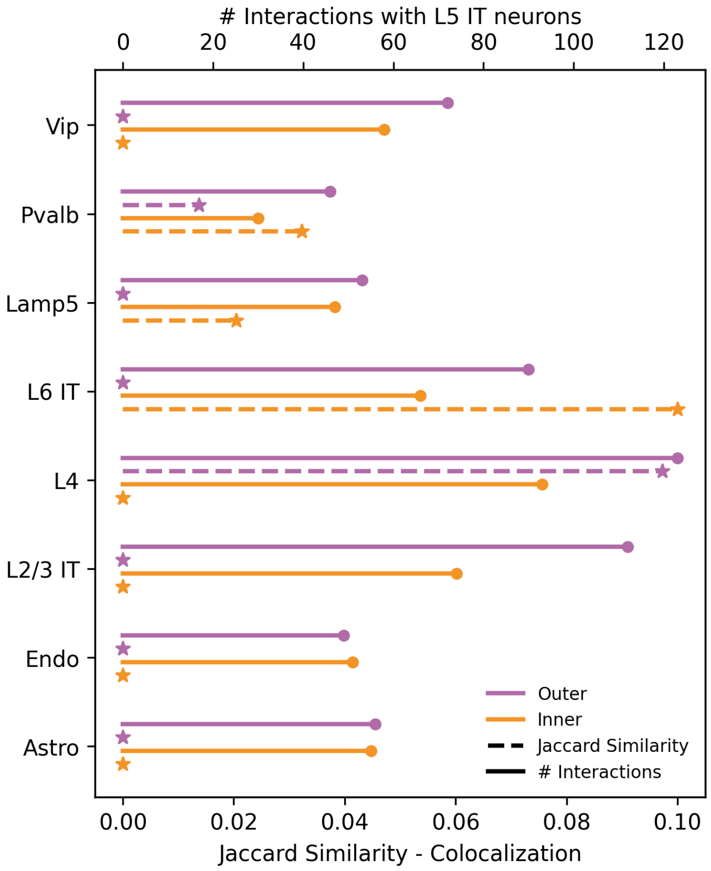


**Figure S11. Comparison between colocalization and LIANA cell-cell communication results in the brain dataset.** This figure presents a comparative analysis of colocalization results and the number of ligand-receptor interactions predicted between inner/outer L5 neurons and cells in the y-axis. Plain line represents the number of significant ligand-receptor interactions identified by LIANA, utilizing the consensus score. Dotted line illustrates the degree of colocalization, measured as the Jaccard similarity index between the spot profiles of L5 neurons and y-axis cells (based on the presence or absence of cells in each spot). For visualization purposes, we selected the extreme cases, those with the highest and lowest levels of colocalization and interaction frequencies.

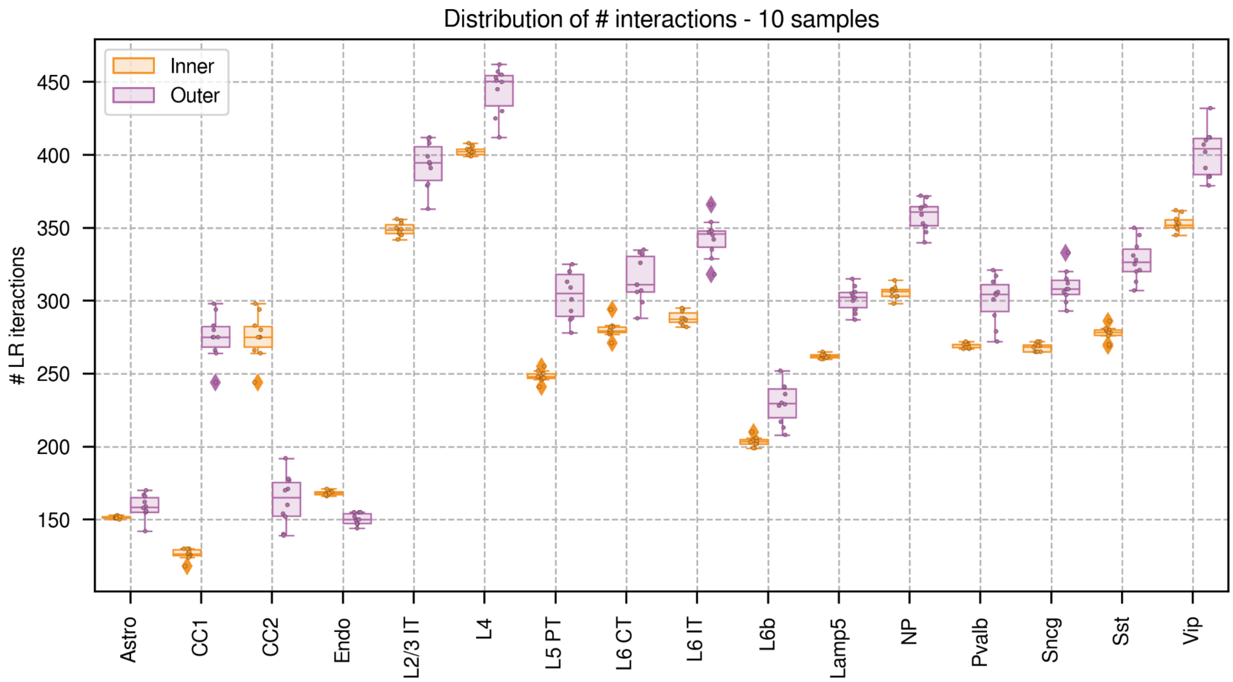


**Figure S12. Subsampling exercise.** This figure illustrates the minimal variability in predicted ligand-receptor interactions counts between L5 IT neurons located in different parts of the L5 layer and the rest of the cells in the dataset, over several subsampling iterations. To ensure a balanced analysis and mitigate biases due to varying neuron quantities in each region, ten unique subsampled datasets were created. Then, we predict the ligand-receptor interactions with CellPhoneDB adjusted for these quantity discrepancies.

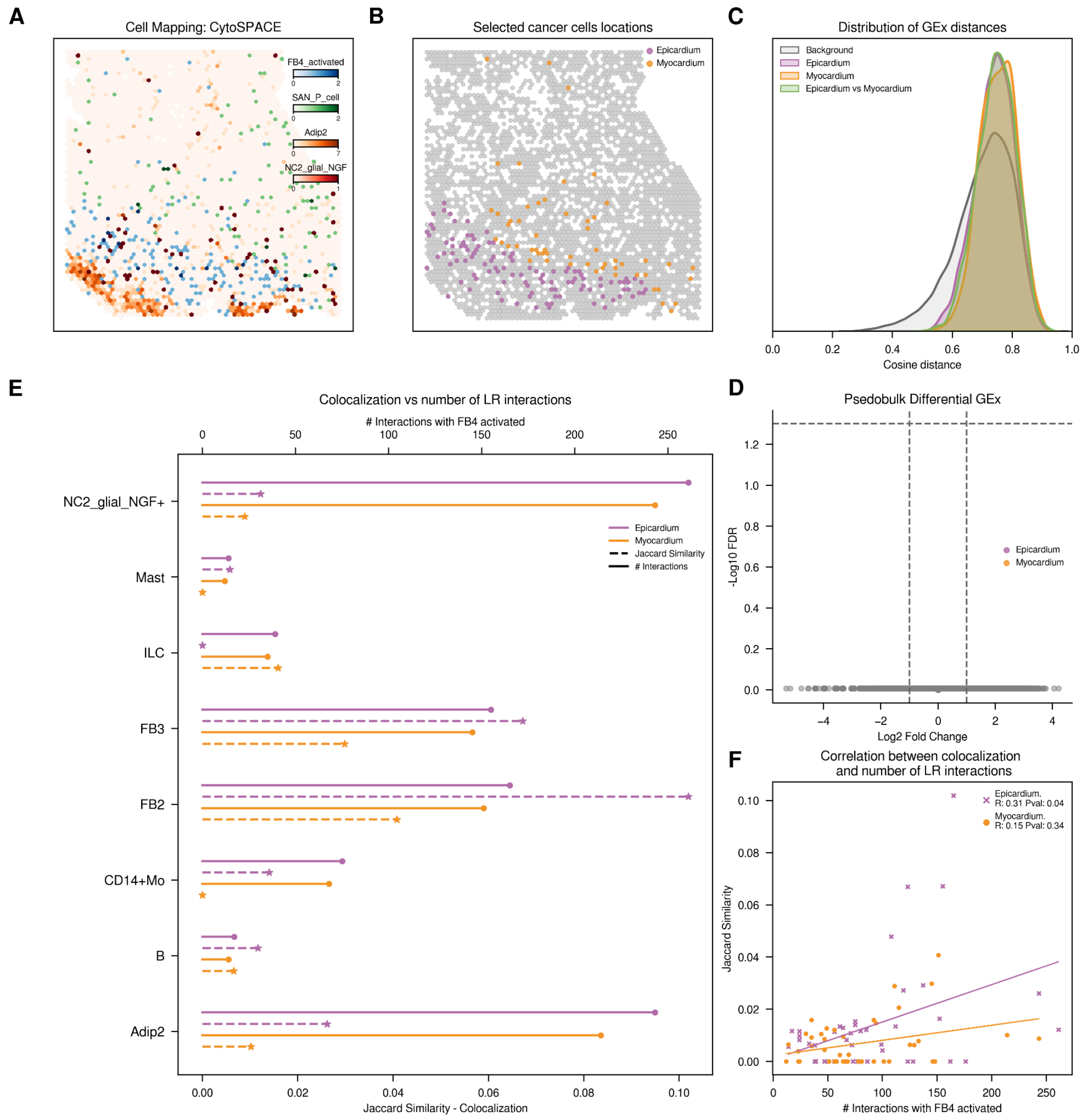


**Figure S13. Exploration of fibroblast (FB4_activated) from SAN slide. A.** Distribution of main cell types across the tissue slide. For visualization purposes, we normalized the CytoSPACE inferred abundances of predominant cell types, with each spatial spot representing the cell type of highest abundance. **B**. Spatial representation of fibroblasts according to their assigned location. **C**. Distribution of cosine distances between gene expression (GEx) profiles. We analyze the cosine distances between gene expression patterns of fibroblasts located within the same or different tissue histological regions. **D**. Volcano plot with Differential Gene Expression results, highlighting no significant upregulated genes in fibroblasts of each histological region. **E**. Comparison of colocalization and CellPhoneDB cell-cell communication results. Plain line (top axis) represents the count of significant ligand-receptor interactions between each of the defined fibroblasts and the y-axis cells. Dotted line (bottom axis) indicates colocalization, measured by the Jaccard similarity index based on the presence or absence of cells within each spot. **F**. Spearman correlation between colocalization and ligand-receptor interaction counts. Color code: Violet fibroblasts in the epicardium, orange fibroblasts in the myocardium.

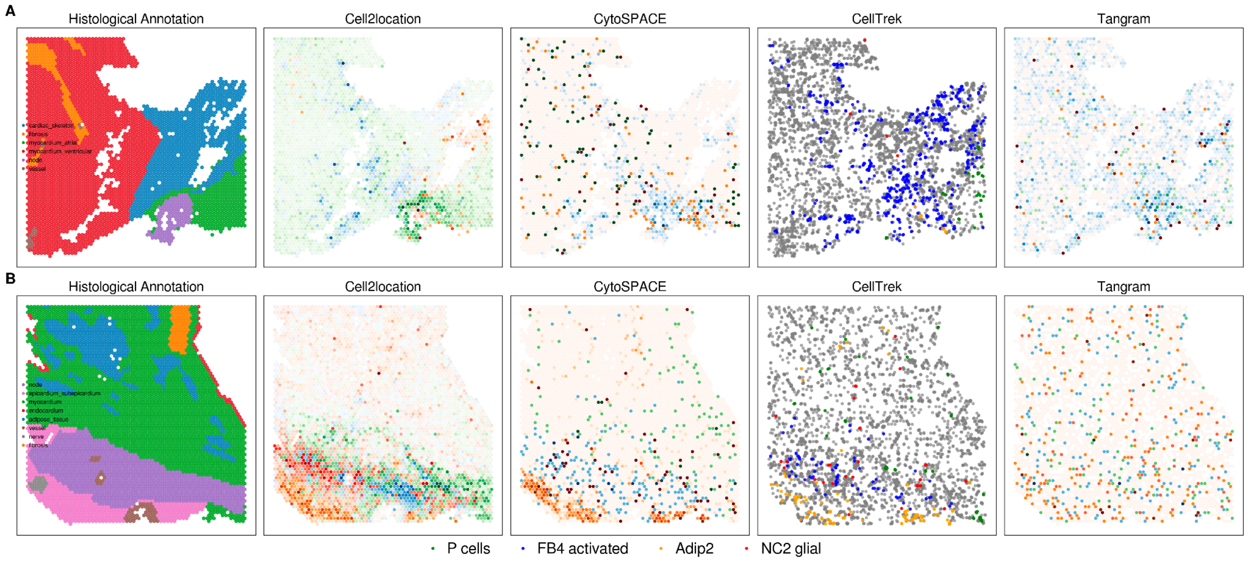


**Figure S14. Methodological comparison for disentangling single-cell spatial resolution for Human Heart Atlas dataset**. We applied four methodologies - Cell2location, CytoSPACE, CellTrek and Tangram - to determine cell positioning across different heart regions (identified as AVN and SAN, and labeled **A**, **B** respectively). Histological annotations from the original authors are included for comparative analysis (Kanemaru et al., 2023), showing partial agreement between histological and Cell2location results (provided by the authors) and the rest of the cell mapping results. Notably, we were unable to map P cells into the histologically annotated nodes. The figure illustrates the normalized abundance of predominant cell types, with each spatial spot representing the cell type of highest abundance for clear visualization. Color coding: green P cells, blue Fibroblast 4 activated, red NC2 glial and orange Adipcites 2.

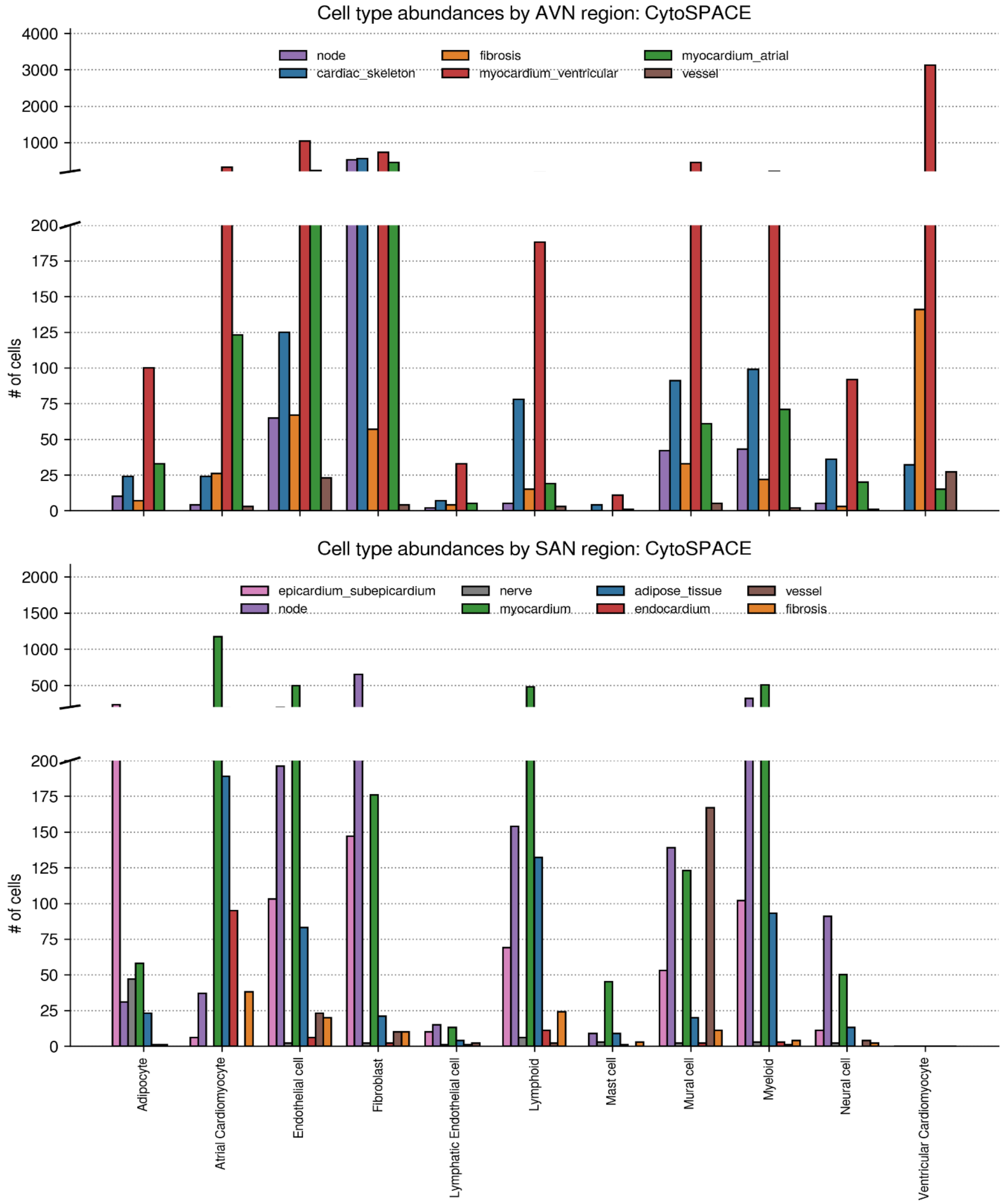


**Figure S15. Variability of cell composition across histological regions in AVN and SAN slides.** Bar plot illustrating the distribution of cell types, determined using CytoSPACE, across the different histological regions defined within AVN and SAN slides. Abundances were calculated by summing the number of cells of each type mapped to specific spots corresponding to the histological regions within the CytoSPACE results. The figure revealed significant compositional variations based on histological region. For instance, the node region on the SAN slide showed a more diverse cell composition compared to the node region in the AVN slide, which was largely dominated by fibroblasts. Each bar represents the count of a specific cell type within histological regions, and distinct colors are assigned to each histological region to facilitate comparison.

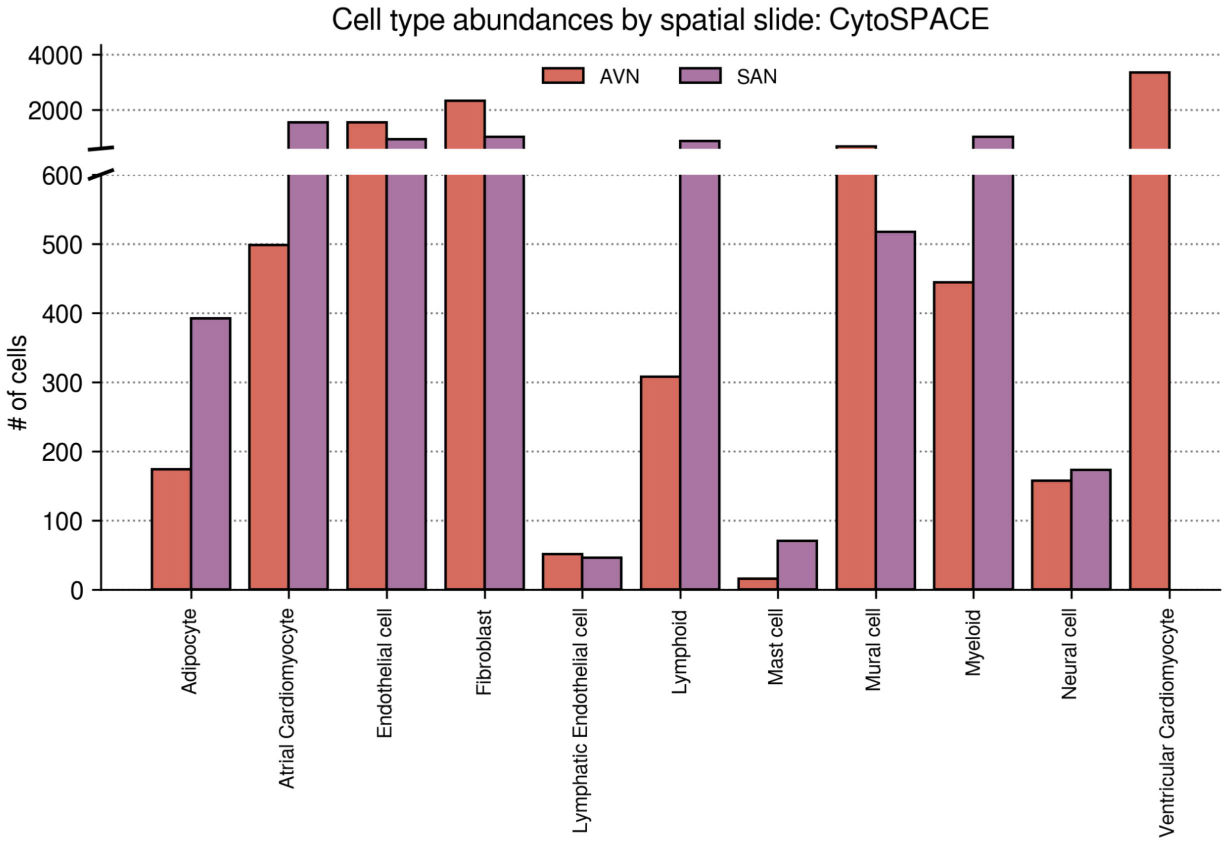


**Figure S16**. **Variability of cell type abundance across heart regions**. Bar plot showing the distribution of cell types, determined using CytoSPACE, across AVN and SAN slides. Abundances were calculated by summing the number of cells of each type mapped to individual spots within the CytoSPACE results. The figure highlights similar cell type proportions across slides, except for ventricular cardiomyocytes exclusive to the AVN slide and a substantial difference in lymphoid cell presence. Each bar represents the count of a specific cell type within a slide, and distinct colors are assigned to each region to facilitate comparison.

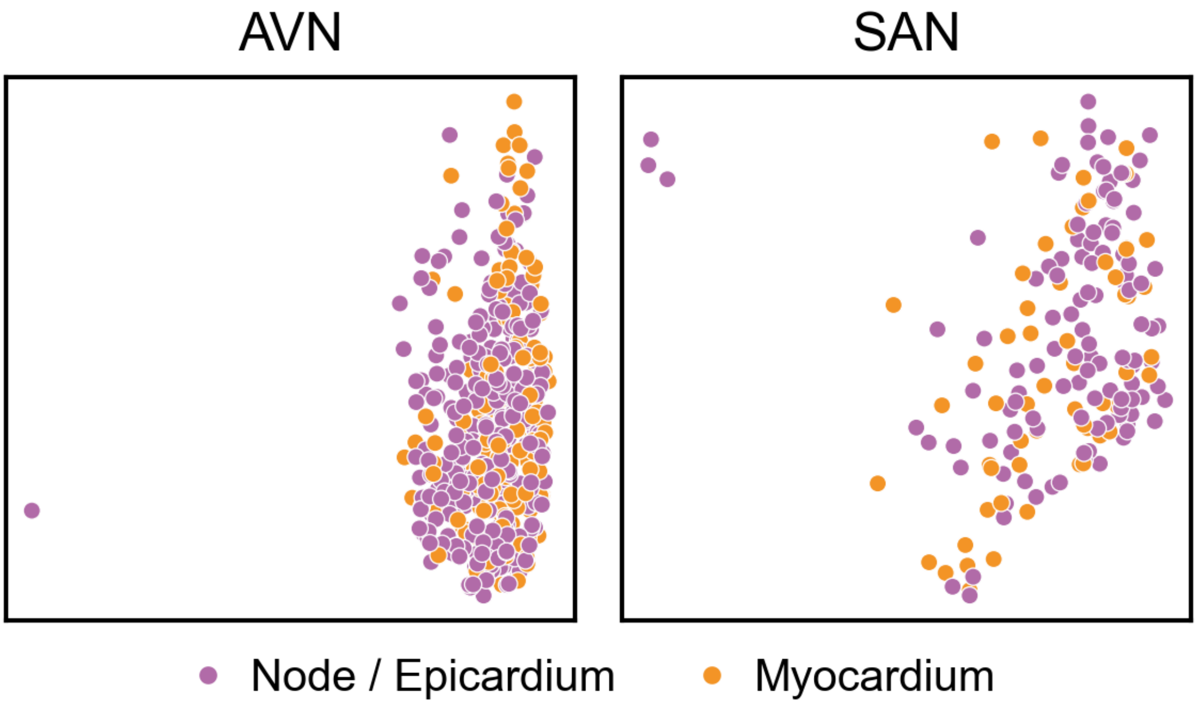


**Figure S17**. **UMAP projection of fibroblast gene expression by location.** Two-dimensional Uniform Manifold Approximation and Projection (UMAP) visualization of the gene expression profiles of fibroblast (FB4_activated) in AVN and SAN regions, aimed to explore the potential unsupervised separation based on the fibroblast assigned location. Each point represents an individual fibroblast, colored according to its assigned location. The distribution appears homogeneously across the UMAP space for both slides, suggesting a lack of differences in the gene expression profiles of fibroblast across different histological regions.

.

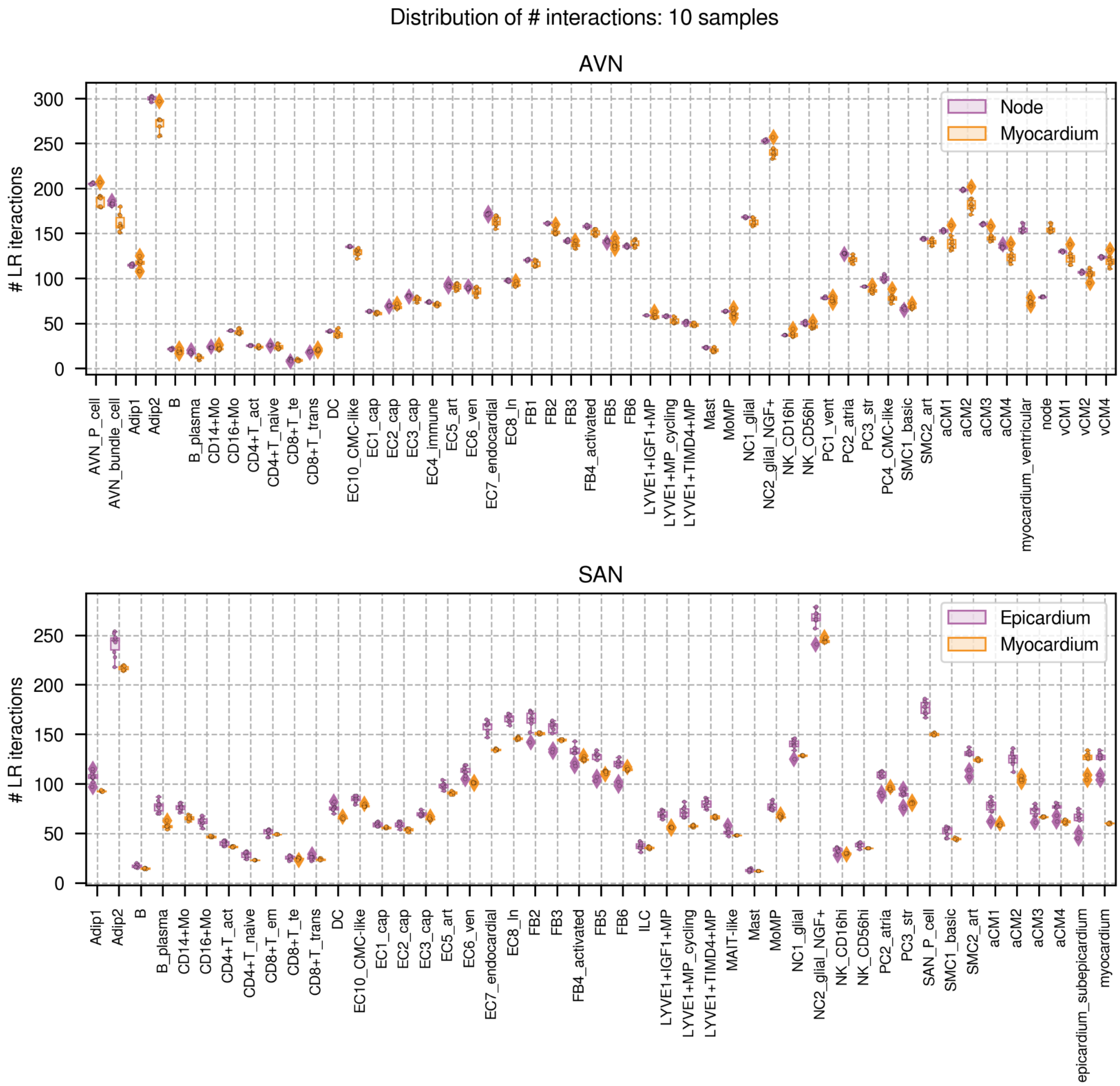


**Figure S18. Subsampling exercise in AVN and SAN slides.** This figure illustrates the minimal variability in predicted ligand-receptor interactions counts between fibroblasts (FB4_activated) located in different histological regions and the rest of the cells in the dataset, over several subsampling iterations. To ensure a balanced analysis and mitigate biases due to varying fibroblast quantities in each region, ten unique subsampled datasets were created. Then, we predict the ligand-receptor interactions with CellPhoneDB adjusted for these quantity discrepancies.

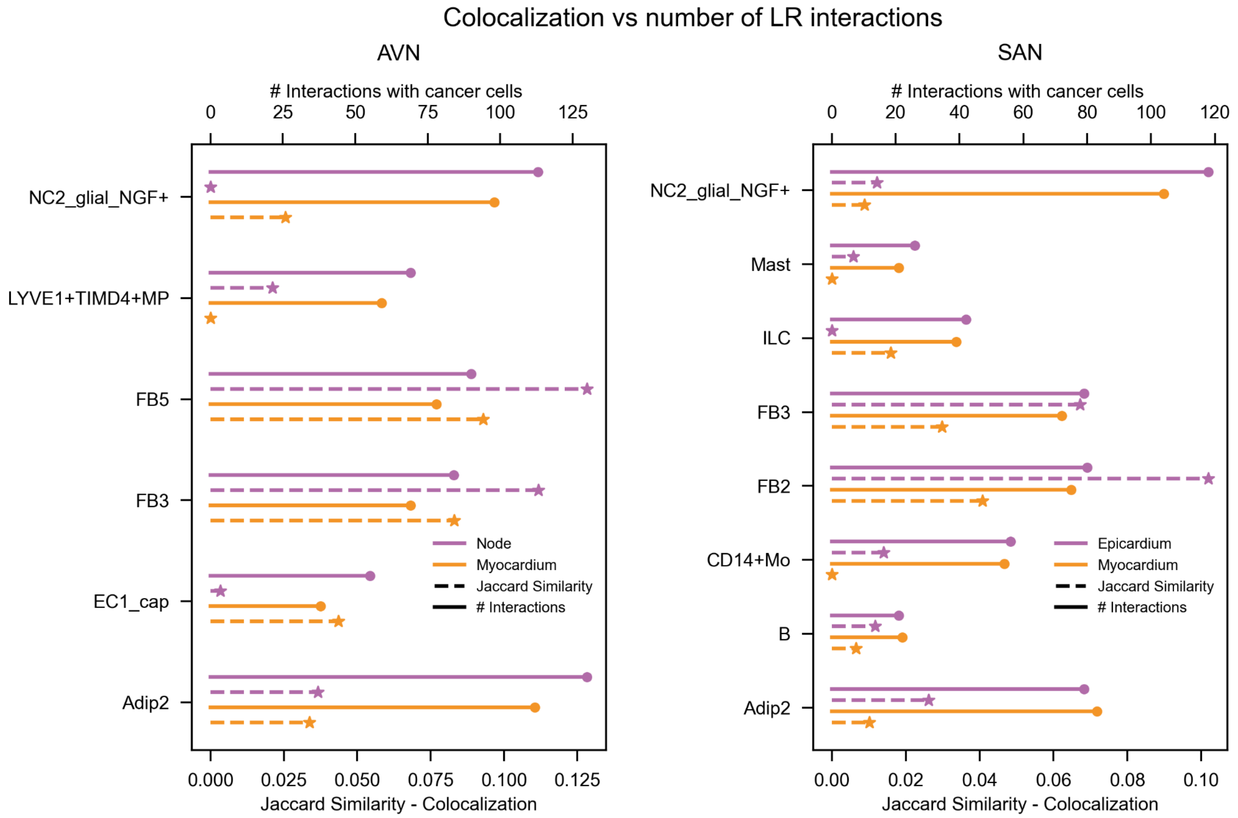


**Figure S19. Comparison between colocalization and LIANA cell-cell communication results in AVN and SAN slides.** This figure presents a comparative analysis of colocalization results and the number of ligand-receptor interactions predicted between histologically-defined fibroblasts and cells in the y-axis. Plain line represents the number of significant ligand-receptor interactions identified by LIANA, utilizing the consensus score. Dotted line illustrates the degree of colocalization, measured as the Jaccard similarity index between the spot profiles of fibroblasts and y-axis cells (based on the presence or absence of cells in each spot). For visualization purposes, we selected the extreme cases, those with the highest and lowest levels of colocalization and interaction frequencies.
